# Migration strategies of a high-latitude breeding songbird (*Setophaga coronata coronata*) revealed using multi-sensor geolocators and stable isotopes

**DOI:** 10.1101/2024.10.09.617415

**Authors:** Stephanie J Szarmach, Johanna K Beam, Mads Moore, Benjamin M Van Doren, Alan Brelsford, David P L Toews

**Author notes:** Corresponding author: Stephanie J. Szarmach.

## Abstract

Seasonal migration allows animals to use habitat where conditions are unfavorable for part of the year but may constrain breeding ranges due to the costs of longer migrations as ranges expand poleward. In species with large ranges, high latitude breeding populations may employ different migratory strategies allowing them to persist far from other core non-breeding areas. The myrtle warbler (*Setophaga coronata coronata*) has two disjunct non-breeding ranges in North and Central America–one along the Gulf Coast and the other on the Pacific. Previous work indirectly linked birds breeding in Alaska with the Pacific non-breeding area, suggesting that high latitude populations evolved a shorter migration route. We directly tested this hypothesis using geolocators measuring both light and atmospheric pressure to track Alaskan myrtle warbler migration in fine detail and inferred non-breeding areas using hydrogen isotopes for a larger sample of birds breeding in Alaska, British Columbia, and Alberta. We found, contrary to expectations, that all geolocator-tracked birds and most birds with stable isotope data migrated to the southeastern United States, while only a small subset of birds (∼5%) likely wintered on the Pacific Coast. We additionally demonstrate the advantages of pressure geolocation for characterizing migratory behavior at a fine scale.

## INTRODUCTION

Seasonal migration allows animals to escape unfavorable environmental conditions and take advantage of spatial heterogeneity in resource availability, predation, and competition by occupying distinct breeding and non-breeding ranges at different times of year [1,2]. In birds, recaptures of banded individuals, genetic data, and, increasingly, remote tracking have shown intraspecific variation in migration routes and non-breeding locations, even among individuals breeding in the same area [3–5]. Linking the breeding and non-breeding grounds of individuals and characterizing migratory connectivity within a species is important both for conservation throughout the full annual cycle and for answering questions about the evolution of migratory traits, as well as the relationship between migration and geographic range.

While migratory behavior permits birds to breed in locations where they could not persist year-round (in the absence of other adaptations like torpor) [2], migration could also constrain species’ breeding ranges, due to the energetic cost of longer migrations resulting from range expansion [6] or an innate predisposition to migrate in a direction that would lead to an inhospitable area from the new breeding location [7,8]. Somewhat counterintuitively, given their breadth of movement, many migratory species are *less* widely distributed than non-migratory species, and occupy smaller ranges than expected based on the presence of suitable habitat [9–12]. For instance, for some migratory species in North America, simulated breeding ranges based on resource availability extend farther north and west than species’ true ranges, suggesting that the actual ranges may be constrained by the distance required to migrate to non-breeding grounds on the Gulf Coast or in Central or South America [12,13]. However, comparative analyses have shown that migratory distance or propensity does not predict range size or range filling across North American breeding birds as a whole [14], and certain migratory species’ breeding ranges do extend into high latitude regions far from their core nonbreeding areas. This raises the question: why do some migratory species breed in high latitude regions where other closely related species with similar habitat preferences do not?

One mechanism that could facilitate breeding range expansion is the evolution of migration routes to new nonbreeding areas located closer to the expanding edge of the range. Models predicting patterns of migratory connectivity have shown that the minimization of energy expenditure (i.e. migration to a closer nonbreeding ground) is an important factor explaining why individuals from different breeding populations winter in different areas [15]. In many species with high migratory connectivity, birds breeding in eastern North America typically migrate to an eastern nonbreeding area, while those breeding farther west winter on a more western nonbreeding ground [16–18]. In some cases, population-level variation in nonbreeding location could arise simply because migrating in the same direction from a more western breeding area results in a corresponding longitudinal shift in nonbreeding area (as demonstrated in translocation experiments) [19,20]. However, in other cases, birds may colonize a novel nonbreeding ground due to a change in migratory orientation. Migration routes to new nonbreeding areas (and, conversely, to new breeding grounds from an existing nonbreeding ground [21]) have evolved in a number of avian species, even within the last fifty years [22,23]. Comparative studies of the relationship between migratory traits and geographic range typically measure migration distance as a straight line between the centroid of a species’ breeding range and the centroid of the non-breeding range, which obscures any within-population variation in migratory direction and distance. Linking the breeding and nonbreeding grounds of individual birds is necessary for understanding how changes in migration direction and nonbreeding area might facilitate breeding range expansion.

The parulid family of wood warblers is a valuable system for studying the relationship between range expansion into the high latitudes of western North America and variation in migratory strategies. Some warblers breeding in the North American boreal forest exhibit a pattern of discordance between their occupied ranges and available habitat [12], and species that migrate farther tend to have smaller ranges [14]. While this suggests that migration constrains range size in this group, a few species do have ranges extending from the Atlantic Coast to Alaska [12]. One of these, the myrtle warbler (*Setophaga coronata coronata*; a subspecies within the yellow-rumped warbler species complex) has two disjunct nonbreeding ranges–one on the Gulf Coast extending to the Atlantic Coast and the other on the Pacific (Figure 1)–making it an ideal candidate for testing the hypothesis that the evolution of alternate migration routes can facilitate range expansion. The myrtle warblers that winter on the Pacific Coast were first described as a subspecies, *S. c. hooveri,* based on their longer wings and tails compared to eastern myrtle warblers, and in this original description it was predicted that they bred in Alaska and British Columbia [24,25]. While *S. c. hooveri* is not considered a distinct subspecies today, analysis by Toews et al. [26] of morphological variation and stable isotopes in myrtle warblers migrating through southwestern British Columbia supported the original prediction that the birds wintering on the Pacific Coast breed at high latitudes in northwestern North America.

**Figure 1.**
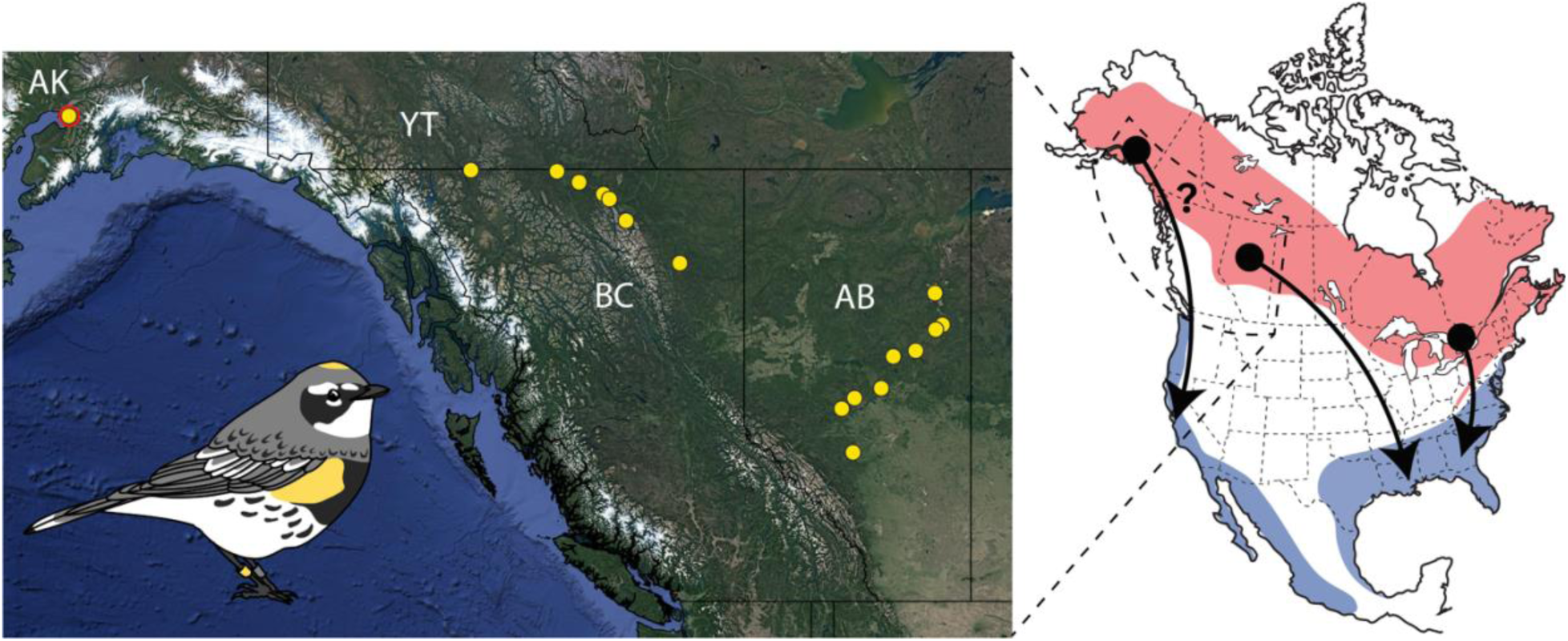
Schematic representing predicted differences in migration route and wintering area for myrtle warblers (*Setophaga coronata coronata*) across their breeding range (Right). The myrtle warbler breeding range is shown in red and wintering range in blue. (Left) Sampling map depicting where geolocators were deployed (Anchorage, AK; point with red outline) and where feather samples were collected for stable isotope analysis (yellow points). Base map: Google Satellite Hybrid (obtained through QuickMapServices QGIS plugin), retrieved 25 January 2024.

However, since Toews et al. [26] sampled myrtle warblers during migration, rather than from the breeding range, it remains unknown whether a migratory divide, a contact zone between populations that migrate to eastern versus western nonbreeding areas, exists. Toews et al. [26] found evidence that most Pacific Coast migrants breed at high latitudes, but further study is needed to determine whether the converse—that most myrtle warblers breeding at high latitudes migrate to the Pacific Coast—is true. Additionally, while stable isotopes from feathers can be used to infer breeding and nonbreeding areas [27,28], these location estimates are often broad due to the natural geographical patterns of isotope distribution, limiting the ability to link precise breeding and non-breeding areas at a fine scale. Geolocator data can provide narrower estimates of nonbreeding areas as well as information about the specific migration paths taken by individuals.

Here, we use multi-sensor geolocators—sensors that quantify both light levels and barometric pressure—to track the migration routes of myrtle warblers breeding in Anchorage, Alaska to test the hypothesis that the evolution of an alternate migration route to a Pacific Coast nonbreeding area facilitated the expansion of the myrtle warbler’s breeding range into high northwestern regions of North America. In addition, we use stable hydrogen isotopes to infer the nonbreeding areas of myrtle warblers breeding in Alaska, northern British Columbia, and Alberta to determine whether a migratory divide is present between birds migrating east to the Gulf Coast and west to the Pacific Coast. We utilize newly developed methods to infer migration trajectories using atmospheric pressure data [29–31], and compare the quality of tracks incorporating pressure versus those based only on traditional light-level data. This pressure data allows us to not only address an evolutionary hypothesis about the relationship between migratory behavior and geographic range, but also to describe migration timing, stopover behavior, and flight altitude—which is currently unknown for this and most other North American migratory bird species.

## METHODS

### Geolocator deployment and sample collection

Between June 1 and June 14, 2022, we captured 55 myrtle warblers using song playback and mist nets in Far North Bicentennial Park and the BLM Campbell Tract, a ≈19 km^2^ forested area located in Anchorage, Alaska (electronic supplementary material, figure S1). All captured birds were banded, measured, and two greater covert feathers were collected for stable isotope analysis. Because of song playback primarily attracting territorial males, all but one captured birds were male. We attached a multi-sensor geolocator (Migrate Tech BARP30Z11-DIP; 0.45g) that measures ambient light intensity, temperature, and air pressure to 30 of these birds using a modified leg-loop harness designed for small songbirds [32]. Geolocators sampled light intensity every minute, recording the maximum intensity every 5 minutes, and recorded pressure and temperature every 20 minutes. In addition to an aluminum USGS band, birds fitted with geolocators were also given a red color band to aid in resighting the following year. To serve as controls when testing for an effect of geolocator on return rate (as, to our knowledge, geolocators have not been used before on this species), 25 birds were processed identically but not given a geolocator and were fitted with a yellow color band to facilitate resighting.

To recapture geolocator-tagged birds, we surveyed the study area from June 1 to June 14, 2023, listening for singing myrtle warblers and playing song audio at each previous capture location and throughout the trails of the park. We resighted and recaptured 6 of 30 birds with geolocators (20%). All units recorded data for the entire deployment period. We resighted 6 of 25 color-banded control birds (24%) and recaptured two. All recaptured birds were confirmed to be the same birds breeding on the respective territory in 2022. We compared return rates of geolocator-tagged birds with control birds using a Fisher’s exact test and found no significant difference in returns (*P* = 0.75).

### Estimation of migration routes and nonbreeding areas from multi-sensor geolocators

We implemented two approaches to estimate locations from the geolocator data to compare the efficacy of the methods. First, we inferred locations using light intensity data alone, largely following the threshold method pipeline described by Lisovski et al. (2020). Then, we modeled migration routes using first atmospheric pressure data alone, and then both pressure and light data together using the R package GeoPressureR [29,30].

For the light-only approach, we first annotated twilight times using the *preprocessLight* function in the R package TwGeos [34] using a threshold of 0.37-1 lux, depending on the tag (electronic supplementary material, table S1 and figure S2). We removed outlier twilight times, which likely resulted from shading, using the *twilightEdit* function in TwGeos and removal criteria of window = 4, outlier.mins = 25, and stationary.mins = 20. In-habitat calibration was performed using the *getElevation* function in the GeoLight R package using a subset of the light data from three weeks after geolocator deployment, when the birds were stationary at their breeding sites (electronic supplementary material, table S1). Visual inspection of the light data using the *LightImage* function in TwGeos showed a long period of increased shading from early July to late August (electronic supplementary material, figure S3). This shading could be due to behavioral changes in the birds during molting: warblers are thought to undergo high-intensity molt, and this loss of flight feathers likely causes them to become more sedentary and shelter under vegetation during this time [35]. A second calibration was performed for this period of increased shading, and the resulting sun elevation angles were used to estimate locations during this time. Shading during migration and on the nonbreeding ground appeared similar to that observed during the first three weeks on the breeding ground, so the sun elevation angle from the first calibration period was used for the remainder of the year. Geographic locations were estimated using the simple threshold method (*coord* function) in GeoLight. We inferred stationary periods and timing of migratory flights using a changepoint model implemented by the *changeLight* function in GeoLight (quantile = 0.9, rise.prob = 0.04, set.prob = 0.05), followed by the *mergeSites2* function (distThreshold = 100, mask = land). We plotted location points for the longest stationary period during winter (“nonbreeding area”), spring migration, and breeding area (final stationary period) using the ggmap [36] and ggplot2 [37] R packages. We removed points between three weeks before and three weeks after the fall and spring equinoxes (1 September 2022—13 October 2022 and 27 February 2023—10 April 2023), because similar day length around the world during this period results in high error for latitude estimates. We did not plot location estimates for fall migration on these maps, because this migration occurred almost entirely during the autumn equinox period for all six birds.

We then inferred migration paths and nonbreeding areas using atmospheric pressure data alone, following the approach outlined in “A User Manual for GeoPressureR” [38]. First, we annotated stationary periods and migratory flights manually in TRAINSET, a web application for labelling time series data [39]. Migratory flights were identified from sudden large drops in pressure, typically >50hPa and lasting at least two hours. Short flights resulting in fine-scale or altitudinal movements were discarded, and extended periods differing in atmospheric pressure within a single stationary period were marked with different elevation labels. We then constructed pressure likelihood maps using the *geopressure_map* function in the GeoPressureR package. Each likelihood map is computed by comparing the pressure recorded by the geolocator with both spatial and temporal variation in pressure extracted from the ERA5-Land surface-level pressure reanalysis dataset [40]. First, a spatial mask is applied to filter out ERA5 grid cells where the pressure at ground level does not match the pressure recorded by the geolocator during a given stationary period. For all remaining grid cells, the normalized mean square error is calculated between the pressure timeseries recorded by the geolocator and the ERA5 timeseries for the grid cell [30].

After initial construction of pressure likelihood maps, we assessed the quality of fit between recorded geolocator data and the ERA5 pressure timeseries by identifying outliers and examining histograms of error (difference between pressure recorded by the geolocator and ERA5) for each stationary period. Stationary periods exhibiting high mismatch between pressure recorded by the tag and ERA5 were probably misclassified during labelling in TRAINSET, and likely represent two different stationary periods that were erroneously combined, or a period when the bird was spending time at different elevations. Pressure timeseries where the bird was spending short times at a different altitude were either marked with an elevation label or discarded, and series where it appeared multiple stationary periods had been combined were split.

Once a good fit between the pressure timeseries recorded by the geolocator and the ERA5 dataset had been achieved for each stationary period, we modeled the full trajectory of each bird’s migration using hidden Markov models implemented in GeoPressureR [29]. A trellis graph was built using the *graph_create* function with a likelihood threshold of 0.99 and groundspeed threshold of 150 km/h. We set a movement model to designate the likelihood of different groundspeeds using a gamma distribution (shape = 7, scale = 7), with a fixed probability for low groundspeeds (<15 km/h) to account for frequent short-distance flights. Using this model, we computed the most likely path taken by each bird using the *graph_most_likely* function and mapped the marginal probability of each stationary period’s location using the *graph_marginal* function to assess uncertainty in location estimates.

Finally, we incorporated both the pressure and light data into a single model in GeoPressureR. Unlike the GeoLight approach, which estimates locations for each pair of twilight estimates and defines stationary periods afterwards based on changes in twilight times, GeoPressureR first aggregates the light data into stationary periods defined by the changes in pressure (i.e. migratory flights) and then estimates the location of each stationary period based on all light readings during that time [30]. For the light analysis, we first estimated twilights from the light data using the *twilight_create* function in GeoPressureR and manually discarded outlier twilight estimates in TRAINSET. The analysis then largely followed the pipeline used for the pressure data alone, outlined above, but in addition to computing the pressure likelihood map, we computed a light likelihood map using the *geolight_map* function. Both the pressure likelihood map and light likelihood map are then used in conjunction with the movement model when generating the most likely trajectory of the bird. We estimated the altitude of the bird during stationary periods and migratory flights using the *plot_pressurepath* function in GeoPressureR, which computes flight altitude using the barometric formula. This function corrects for natural fluctuations in barometric pressure using the ERA5 pressure data, and uses the locations estimated for the stationary periods preceding and following the flight to retrieve the ground level pressure and temperature for the region over which the bird is flying.

After migration tracks and nonbreeding areas had been estimated using the three different methods (light data alone in GeoLight, pressure data alone in GeoPressureR, and both pressure and light in GeoPressureR), for each bird we plotted the products together on the same map to compare the results. We plotted the nonbreeding areas estimated using each method using the ggmap and ggplot2 R packages, plotting the likelihood rasters for the longest winter stationary period from the GeoPressureR analysis using *geom_spatraster* and the location point estimates generated in GeoLight for wintering period using *geom_point*. We then compared migration routes estimated using the different methods by plotting the tracks generated from GeoPressureR and the location points estimated using GeoLight on the same base map.

We visualized differences in migration timing between individuals using the vistime R package to create timeline plots [41]. We compared migration timing characteristics (duration, number of flights, length of flights, length of stationary periods) between fall and spring using t-tests or Wilcoxon rank sum tests if the data was not normal, implemented using the R packages stats or exactRankTests, respectively [42,43].

### Estimation of nonbreeding areas from stable isotopes

We inferred nonbreeding areas in a larger sample of myrtle warblers breeding across northwestern North America using stable hydrogen isotope analysis. The ratio of hydrogen isotopes (deuterium, ^2^H, to protium, ^1^H, or δ^2^H) in a feather can be used to infer the location of a bird at a certain time because of a strong correlation between the δ^2^H of a feather and the δ^2^H of precipitation where the feather was grown [27]. Adult myrtle warblers undergo both a complete molt on the breeding ground and a partial molt on the nonbreeding ground, so both locations can be inferred using stable isotopes from different generations of feathers on the same bird [26,44,45]. We measured the stable hydrogen ratio (δ^2^H) in greater covert feathers grown during the pre-alternate molt (i.e. on nonbreeding grounds) for 167 myrtle warblers breeding in Alaska (*n* = 46), British Columbia (*n* = 54), and Alberta (*n* = 67; Figure 1). In addition, we assessed δ^2^H in feathers grown during the pre-basic molt (i.e. on breeding grounds) for 24 myrtle warblers breeding in Alaska (*n* = 12) and British Columbia (*n* = 12) to assess how well the location of feather origin inferred using stable isotopes matched the known breeding location. Alternate and basic greater covert feathers were identified based on feather wear, as alternate feathers grown more recently on the nonbreeding ground are darker black, with bright white tips, and basic feathers are duller grey with more worn and yellowed white tips (electronic supplementary material, figure S4). Feather sample preparation and hydrogen pyrolysis were conducted at the Cornell University Stable Isotope Laboratory. The hydrogen isotope ratio δ^2^H was determined by comparing the ratio of deuterium to protium in the feather sample to the ratio in established keratin standards corrected against the Vienna Standard Mean Ocean Water (VSMOW; electronic supplementary material, table S2).

We assessed the expected distributions of isotope values for the eastern and western wintering grounds by sampling the precipitation δ^2^H at random points within each range from a precipitation isoscape model produced using IsoMap (Bowen et al. 2014). Yellow-rumped warbler range shapefiles were downloaded from BirdLife International. Because these shapefiles combine the ranges of all subspecies, and higher concentrations of wintering myrtle warblers are found in a subset of the range, we modified the shapefiles in QGIS to better represent the common wintering range of myrtle warblers based on eBird observations for myrtle warblers in January-February and the eBird Status and Trends abundance map for yellow-rumped warblers (electronic supplementary material, figure S5; Fink et al. 2023). After sampling precipitation δ^2^H at 1000 random points generated inside each “common” wintering area shapefile, we found that the expected distribution of δ^2^H is significantly different between the eastern and western wintering areas (Wilcoxon rank sum test *W*=965705, *P*<2.2e^-16^; electronic supplementary material, figure S6). Based on this result, we expect feathers from birds that wintered on the Pacific Coast to have lower δ^2^H values.

To test the hypothesis that a migratory divide exists between myrtle warblers breeding in Alaska and those breeding further east, we performed a linear regression in R with longitude of the breeding site as the predictor and δ^2^H of the alternate feather as the dependent variable. We also performed an ANOVA followed by Tukey’s HSD test to identify which sampling sites exhibited significantly lower δ^2^H values suggestive of Pacific Coast wintering. To assess whether basic feathers (grown on the previous year’s breeding ground) exhibited expected differences in δ^2^H based on the birds’ known breeding locations, we performed a Wilcoxon rank sum test in R to determine if there is a significant difference in δ^2^H between basic feathers collected in Alaska versus British Columbia.

We estimated the most likely area of geographical origin for each feather sample using the R package assignR [48]. First, we calibrated a global growing season precipitation hydrogen isoscape (“GlobalPrecipGS”) using known-origin isotope data from feather samples of 19 parulid warbler species (313 samples; Hobson and Wassenaar 1997, Hobson et al. 2012, Hobson and Koehler 2015) and the *calRaster* function in assignR. This regression model generated the transfer function: δ^2^H_feather_ = 0.71* δ^2^H_precipitation_ – 17.12 (R^2^ = 0.81; electronic supplementary material, figure S7). We used the *pdRaster* function in assignR to produce calibrated isoscapes for each analyzed feather sample and plotted posterior probability density maps using the R packages ggmap and ggplot2. To assess whether each alternate feather was more likely to have been grown on the eastern or western wintering ground, we used the *oddsRatio* function in assignR to compare the posterior probabilities within the polygon representing the “common” western area vs. the polygon for the “common” eastern area. When estimating the odds that a sample originated from the western wintering ground rather than the eastern wintering ground, an odds ratio higher than 0.75 (the ratio of the two areas) suggested a higher likelihood that the feather was grown on the West Coast.

## RESULTS

All geolocator-tracked myrtle warblers wintered in the southeastern United States, ranging from southern Texas to northern South Carolina (Figure 2). The nonbreeding areas estimated from light-level data alone using the GeoLight R package largely overlapped with those inferred in GeoPressureR using either light level alone, atmospheric pressure alone, or the full model combining light, pressure, and groundspeed probability (Figure 2b, electronic supplementary material, figures S8 and S9). Point estimates of either the highest likelihood nonbreeding location from the GeoPressureR analysis or the median latitude and longitude of points estimated for the wintering period in the GeoLight analysis were highly similar (electronic supplementary material, table S3), but with a greater difference between the two methods for latitude (mean difference: 1.43°, range: 0.05–4.53°) than longitude (mean difference: 0.44°, range: 0.01–0.44°).

**Figure 2.**
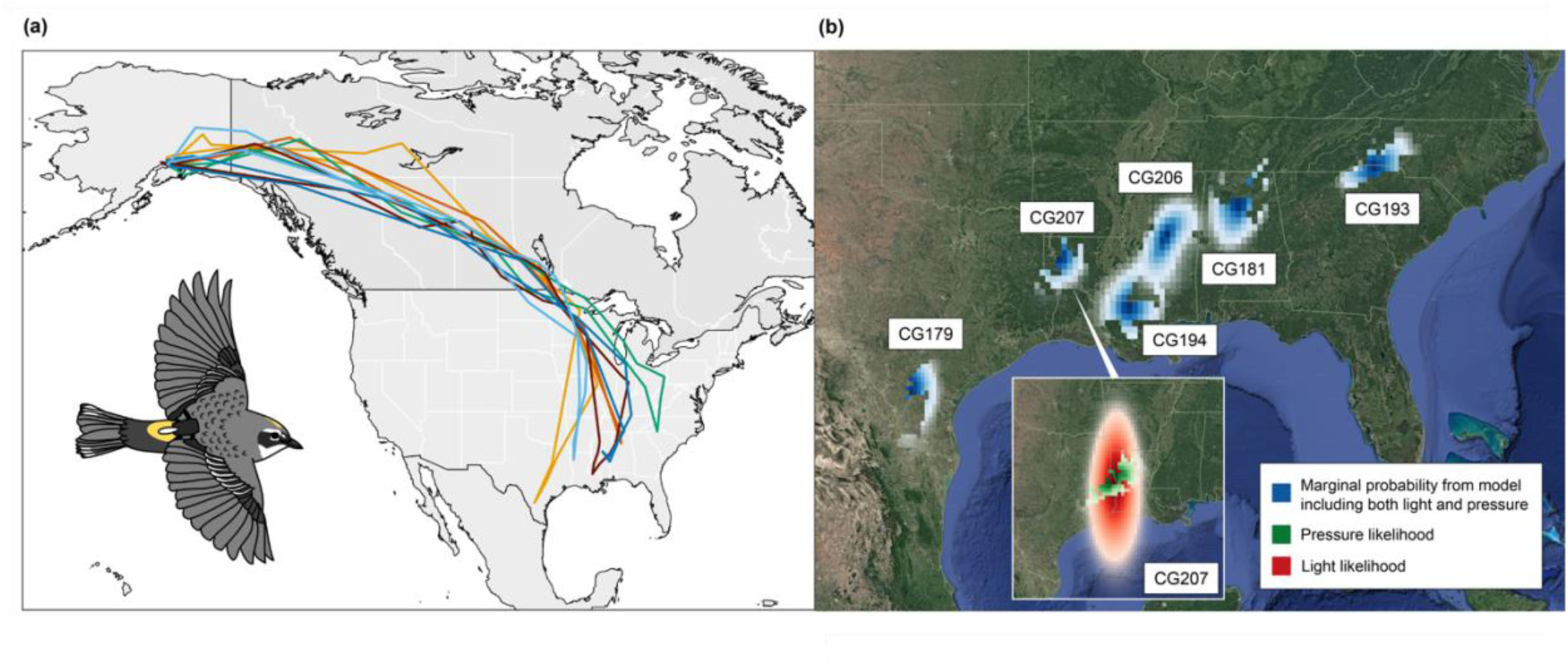
Migration routes and wintering areas of geolocator-tracked myrtle warblers breeding in Alaska. (a) Full-year tracks inferred using combined light and atmospheric pressure data for six myrtle warblers. Each bird’s track is represented by a different color. (b) Probability of wintering location for each bird produced by modeling the birds’ trajectories in GeoPressureR using both light and pressure data collected over the longest winter stationary period (blue grid cells; darker color represents higher probability). The wintering area of each bird is labeled with its geolocator ID. Inset shows likelihood of wintering location derived from light level data alone (red) and pressure data alone (green) for one bird. Base map: Google Satellite Hybrid (obtained through QuickMapServices QGIS plugin), retrieved 29 March 2024.

All six birds followed largely similar routes from Alaska through Yukon and the Interior Plains of Canada, before their paths diverged after passing west of Lake Superior as each bird traveled to a different nonbreeding location in the southeastern United States (Figure 2a; electronic supplementary material, figures S10, S11, S12). The paths followed for fall and spring migration by an individual bird were also similar. Instances where the tracks of individual birds seemed to diverge from the common path through Canada are likely attributable to error in the location estimates, as the likelihood surfaces for these periods were usually broad and also included the area that the other birds passed through. The migration paths inferred from light data alone using the GeoLight package, from pressure data alone using GeoPressureR, and from the combination of light and pressure data were similar for each bird (electronic supplementary material, figure S13). There were some cases where the routes estimated using different methods diverged, likely because of uncertainty in location estimates for certain time periods due to either shading error in the light data or broad pressure likelihood surfaces.

The atmospheric pressure data allowed us to characterize migratory timing in greater detail than using light-level data alone, because each migratory flight could be identified from sharp drops in pressure. By comparing the timing of these drops in pressure to the light levels recorded by the geolocator, we found that almost all migratory flights took place in periods of darkness (electronic supplementary material, figure S14), with the exception of some short flights that occurred during the day and very long flights that extended from night into the morning hours. The average length of a migratory flight was 6.75 hours, and the longest flight was 17.7 hours.

The birds departed their breeding grounds in Anchorage between 28 August and 5 September and arrived on their nonbreeding grounds between 29 October and 4 December (electronic supplementary material, table S4 and figure S15). The birds spent between 89 and 158 days on their nonbreeding grounds, which we defined as the location of the longest winter stationary period (mean = 129, sd = 26.9), and departed for spring migration between 25 February and 9 April. They arrived back to the breeding grounds between 6 May and 19 May. The mean duration of spring migration (47 days) was significantly shorter than that of fall migration (77 days; *W*=33, *p*=0.015), and spring migration was shorter than fall migration for all but one bird (electronic supplementary material, figure S16 and table S4). The birds completed spring migration in fewer flights (mean fall flights = 18, mean spring flights = 15, *W* = 32.5, *p* = 0.022; electronic supplementary material, figure S17 and table S4), and the average length of these flights was longer in spring than fall (mean flight length per bird in fall = 5.8 hours, in spring = 8.0 hours; *t* = –4.23, df = 6.7, *p* = 0.004; electronic supplementary material, figure S18 and table S4). The length of stationary periods (“stopovers”) between migratory flights was significantly greater in fall than spring (mean total stationary days in fall = 73, spring = 42, *W* = 33, *p* = 0.015; electronic supplementary material, figure S19 and table S4). When comparing estimates of migration timing generated using pressure data alone in GeoPressureR to those from light-level data alone in GeoLight, both methods identified large shifts in migratory behavior occurring at similar times (i.e. departure from the breeding ground, arrival on the nonbreeding ground; electronic supplementary material, figure S20). However, GeoLight often split long stationary periods identified from the pressure data (time on breeding ground and nonbreeding ground) into multiple shorter stationary periods and merged shorter stopover periods during migration.

From the atmospheric pressure data collected by the geolocators, we were also able to characterize the flight altitudes of migrating myrtle warblers (electronic supplementary material, figures S21 and S22). The maximum altitude attained by any individual was 2903m above sea level. The median flight altitude for each bird over the full year ranged from 721m to 919m a.s.l. (average = 839m a.s.l.). Though two birds flew at significantly lower altitudes in spring compared to fall (electronic supplementary material, figure S23), overall, there was no significant difference in median flight altitude between seasons across all birds (*t* = 1.25, *df* = 6.1, *p* = 0.26). The pressure data also allowed us to characterize finer scale vertical movements, such cyclical changes in pressure corresponding to periods of daylight and darkness when the birds were likely foraging and roosting at different elevations, and altitudinal changes occurring within a single flight (electronic supplementary material, figure S24).

Stable isotope values (∂^2^H) of basic feathers (expected to have been grown on the previous year’s breeding ground) significantly differed between samples collected in Alaska versus British Columbia (*W* = 117, *p* = 0.008; electronic supplementary material, figure S25), with BC samples exhibiting lower ∂^2^H (mean ∂^2^H = –128) than AK samples (mean ∂^2^H = –119). The posterior probability density maps of most basic feather samples included the sampling location (breeding location in 2022) within the region of likely origin (electronic supplementary material, figures S26 and S27), but for about half of the Anchorage-breeding birds, the posterior probability was low for the sampling site (*n* = 5).

There was a significant relationship between stable hydrogen isotope values (∂^2^H) in alternate feathers and breeding site longitude (β = 0.425, *df* = 165, *t* = 7.58, *p* = 2.3e^-12^, R^2^ = 0.254; Figure 4). When comparing the posterior probabilities of feather origin from the West Coast nonbreeding ground to the East Coast nonbreeding ground, eight samples had odds ratios greater than 0.75 (the ratio of the nonbreeding ground areas), suggesting that these birds were more likely to have wintered on the Pacific Coast (Figure 4; electronic supplementary material, figure S28). Five of the birds with high likelihood of wintering on the Pacific Coast had bred in Alaska, two in British Columbia, and one in Alberta. The remaining 159 birds had higher odds of wintering on the East Coast nonbreeding area, although the posterior probability density maps of many of these birds showed high likelihood regions within the Pacific Coast nonbreeding area.

**Figure 3.**
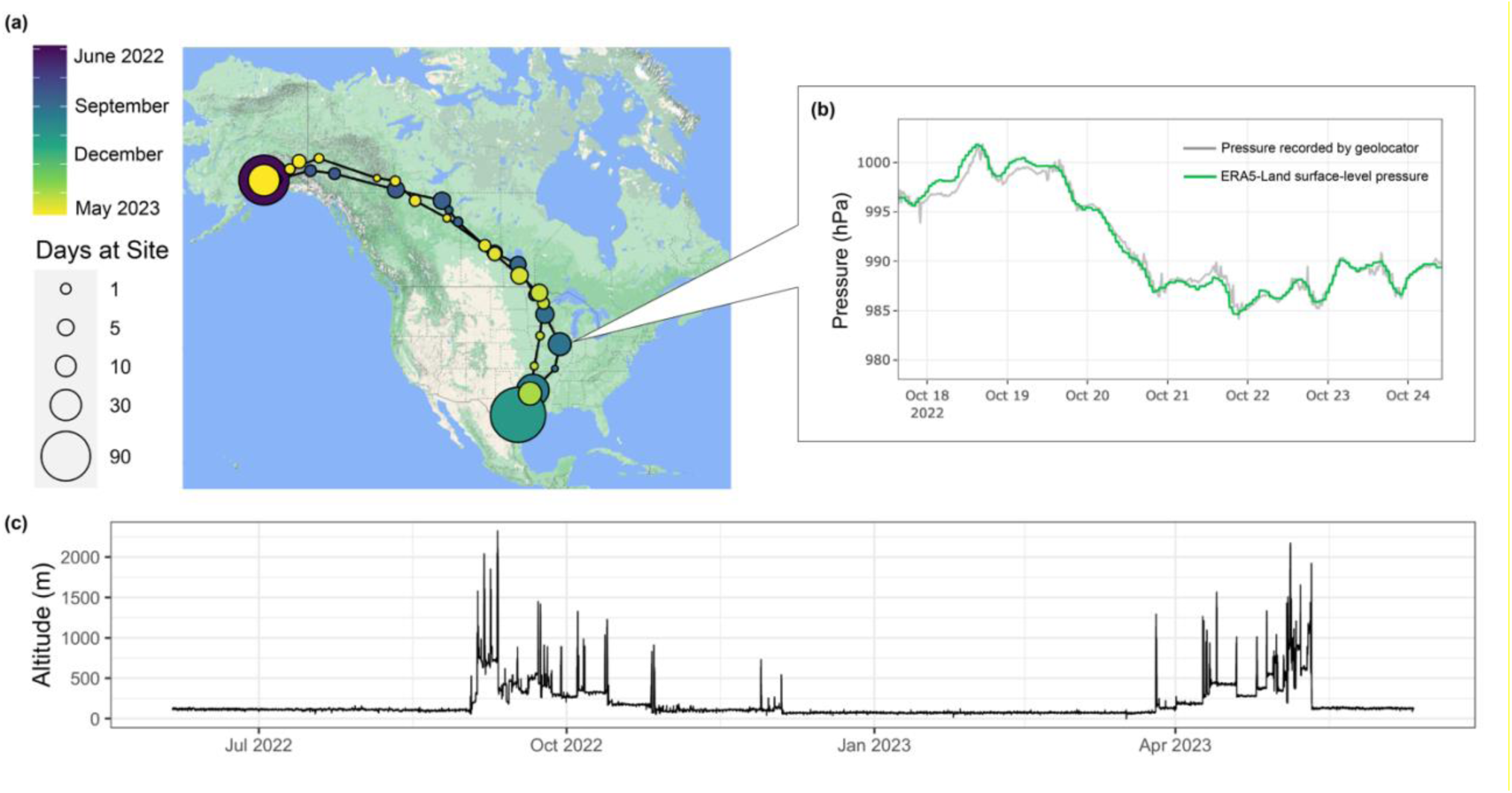
Detailed information about the migration of one geolocator-tracked myrtle warbler gained using atmospheric pressure measurements. (a) Year-long migration route inferred from pressure data alone using the GeoPressureR package. The size of each circle represents the amount of time spent at a site, and the color represents time of year (fall migration: blue, spring migration: yellow). (b) Atmospheric pressure (hPa) measured over time by the geolocator (grey) during a stationary period versus that recorded in the ERA5-Land surface level pressure data set (green) for the same period at the inferred stopover location. A close fit between the two plots provides strong evidence that the inferred stopover location is accurate. (c) Altitude of the bird over the year estimated from the geolocator’s pressure measurements. Sudden sharp peaks in altitude occur due to migratory flights. Map data ©2024 Google, INEGI.

**Figure 4.**
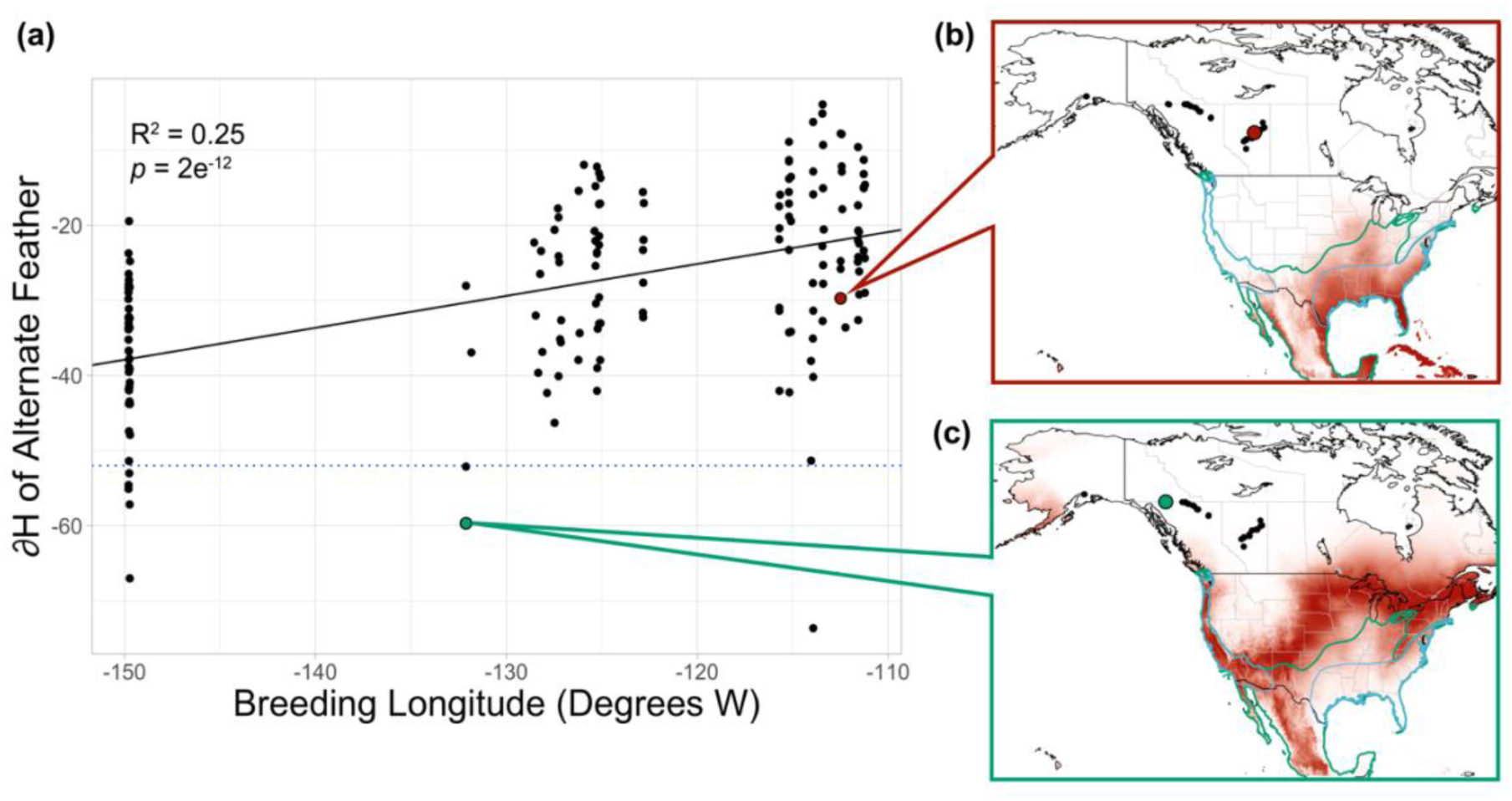
Stable hydrogen isotope analysis of alternate covert feathers sampled from myrtle warblers breeding in Alaska, British Columbia, and Alberta. (a) There is a significant positive relationship between hydrogen isotope ratio (∂^2^H_f_) and breeding longitude (*p* = 2e10^-12^). Points above the dotted blue line have ∂^2^H_f_ values more suggestive of East Coast wintering, while those below have a higher likelihood of having been grown on the West Coast nonbreeding ground, according to an odds ratio test. Posterior probability maps are shown for (b) one individual that likely wintered on the East Coast and (c) one individual with a higher probability of wintering on the West Coast. Darker red areas of the map represent regions of higher probability of origin for the feather sample. The green outline surrounds the full yellow-rumped warbler wintering area, and the blue outline surround the East Coast and West Coast wintering areas where myrtle warblers are most common, according to eBird abundance data. Small black points on the maps indicate all locations where alternate feathers were sampled, and larger red (b) or green (c) points show the location of the highlighted sample.

## DISCUSSION

We characterized the migration routes, migration timing, flight altitudes and wintering locations of six myrtle warblers in fine detail using multi-sensor geolocators, and estimated wintering areas for 167 individuals using stable hydrogen isotope analysis. Additionally, we compared methods of inferring migration routes and timing using traditional light-level data versus atmospheric pressure. Using pressure data allowed us to not only characterize migration routes with less error at finer spatial and temporal scales, but also revealed novel details about migratory behavior, such as flight altitude. We found that, contrary to expectations, all geolocator-tracked birds migrated to the southeastern United States, rather than undergoing a shorter migration to the Pacific Coast wintering ground, as we had originally predicted. In the larger sample of birds for which we estimated wintering areas using stable isotope analysis, most birds (95%) likely wintered in the southeast, while a small subset (5%) showed a higher likelihood of wintering on the Pacific Coast.

### (a) Geolocator and isotope data show that most “hooveri”-type myrtle warblers do not winter on the Pacific Coast

Previous work suggested that myrtle warblers breeding in northwestern North America— once categorized as the *S. c. hooveri* subspecies on the basis of morphology (longer wings and tails in western birds)—wintered on the Pacific Coast [24,26,50]. These morphological differences have previously been associated with variation in migratory behavior in the yellow-rumped warbler species complex [51]. While we observed in our sample that wing and tail length significantly increased in birds breeding farther northwest (electronic supplementary material, figures S29 and S30), with the largest birds in Anchorage, we found that almost all of these birds wintered in the southeastern U.S., regardless of size. The sizes of the geolocator-tracked birds fell throughout the full size distribution of birds banded in Anchorage (electronic supplementary material, figure S31a), so the fact that all of these birds wintered on the Gulf Coast was not the result of a size bias where only smaller birds were tracked. In the larger sample of birds with stable isotope data, the eight birds that most likely wintered on the Pacific Coast all exhibited long wings and tails within the *“hooveri”* range, but almost half the birds that likely wintered on the east coast also exhibited this large size (electronic supplementary material, figure S31b). Thus, while previous studies and our isotope work show that myrtle warblers wintering on the Pacific Coast exhibit the larger *“hooveri”* phenotype, size does not predict whether these birds migrate west or east, and only a small subset of the *hooveri* birds may take an alternate migration route to the Pacific Coast.

The Pacific Coast nonbreeding grounds and northwestern breeding grounds were previously linked using stable hydrogen isotope analysis of feathers collected from myrtle warblers migrating through Vancouver [26]. Isotope values suggested that these birds migrating down the Pacific Flyway bred in Alaska, Yukon, or northern British Columbia and wintered on the Pacific Coast, but—because only one migration site was sampled—this study could not characterize migration direction across the northwestern breeding range and determine if a migratory divide was present. By sampling feathers across a transect of breeding sites from Alaska to northern British Columbia and Alberta, we determined that there is no clear migratory divide between western birds migrating to the Pacific Coast and birds farther east migrating to the Gulf and Atlantic Coasts. Instead, based on a low percentage of samples with low isotope values suggestive of Pacific Coast wintering, it seems that a subset of birds in these areas may migrate to the Pacific Coast, with the proportion of birds that do so increasing in breeding populations farther west. This pattern is similar to what has been observed in Eurasian blackcaps, where birds that take the recently evolved, alternate migration route to winter in Great Britain originate from across the breeding range, representing a low frequency phenotype in multiple breeding populations [4].

Assigning geographic locations of origin using stable isotopes is limited by low spatial resolution due to the broad distribution of isotopes across North America, but integrating multiple types of data (e.g. tracking, banding, community science observations) with stable isotope analysis can help to both corroborate conclusions drawn from stable isotopes and constrain the geographic assignment of samples [52,53]. To validate our isotopic analysis, we analyzed a subset of basic covert feathers (expected to have been grown on the previous year’s breeding grounds based on previous work; [44,54,55] and compared the geographic assignment from the isotope data to the known breeding site where the samples were collected. The region of high likelihood of origin inferred using isotopes overlapped the sampling location for all birds breeding in British Columbia, but only 58% of the samples from Anchorage. All individuals whose isotopic assignments did not match the sample location were second-year birds, so this discrepancy could reflect natal dispersal. Alternatively, because few known-origin feathers from Alaska were available to calibrate the precipitation to feather hydrogen transfer function, this difference could reflect uncertainty in the isoscape for this region [28]. That said, since more known-origin samples are available for the continental United States, this uncertainty is less of a concern for inferring unknown wintering areas.

To validate the isotopic assignment of wintering areas, we compared the posterior probability maps to the geolocator tracks and found that for all individuals with both geolocator and isotope data (*n*=4) the wintering site inferred from geolocators fell within the region of high isotopic likelihood (electronic supplementary material, figure S32). We were not able to validate samples with high isotopic likelihood of wintering on the Pacific Coast in this way, since no geolocator-tracked birds wintered there. While geographic likelihood assignment surfaces generated from hydrogen isotopes are broad—and the same isotope ratio can suggest origin from the Pacific Coast, upper Midwest, or New England—based on eBird observation data myrtle warblers are much less likely to winter in the northeast or upper Midwest. From USGS banding records where myrtle warblers banded in Alaska were re-encountered in the eastern U.S. (*n*=8), the birds found during winter (January-March) were located in Louisiana, Arkansas, and Texas, while birds encountered farther north (Iowa, Wisconsin, and Minnesota) were recovered during fall or spring migration [56]. However, we cannot fully rule out the possibility that the birds with low ∂^2^H were among the few myrtle warblers that winter in the northeast.

It is possible that some feather samples could have lower ∂^2^H because the birds were molting while on spring migration, rather than on the wintering grounds. In Portland, Oregon, Gaddis (2011) found that 40% of myrtle warblers migrating through the area in spring showed prealternate molt, although no studies have examined whether such molt migration also occurs in birds migrating southeast. We typically collected the innermost alternate greater covert feather, which would have been grown earliest during the molt (in myrtle warblers usually beginning in February; Hunt and Flaspohler 2020). However, we cannot completely rule out that this isotopic signature is the result of molting during migration, and more work is needed to characterize temporal patterns of molt in yellow-rumped warblers migrating through the Mississippi Flyway.

Given the low number of birds with a high likelihood of Pacific Coast wintering that we identified in our sample, future work is needed to track myrtle warblers from other parts of the northwestern breeding range or directly from the Pacific Coast wintering grounds to determine where most of the Pacific Coast-wintering birds breed. While Toews et al. (2017) did find areas of high probability of origin in Alaska and northern British Columbia, much of the region of high likelihood encompassed Yukon and the western Northwest Territories (i.e. farther north than where our geolocator birds were tagged). It is possible that a greater proportion of birds breeding in those regions, which were not sampled for the present study, migrate to the Pacific Coast. Alternatively, myrtle warblers breeding in southeast Alaska may be more likely to winter on the Pacific Coast, undergoing a much shorter migration. Though Toews et al. (2017) did not find a high likelihood of breeding in this area for the birds migrating through Vancouver, USGS banding records report that one bird banded in August on the Alaskan coast northwest of Glacier Bay was recovered in California in November [56]. An alternative approach would be to deploy geolocators on myrtle warblers during winter on the Pacific Coast nonbreeding ground. This approach was successful for identifying the breeding areas of Eurasian blackcaps wintering in Great Britain, when deploying geolocators on breeding blackcaps in continental Europe identified only a few instances of birds migrating to Britain [4]. However, nonbreeding site fidelity is not well studied in myrtle warblers, and may be lower than breeding site fidelity [57], which could result in lower return rates and fewer recovered geolocator tracks.

### (b) Most myrtle warblers breeding in northwestern North America follow a similar migration route to other songbird species that may retrace historical range expansion

Though few studies have tracked the migrations of songbirds breeding in northwestern North America, the route followed by myrtle warblers in this study was similar to routes tracked in other songbirds breeding in Alaska, including tree swallows (*Tachycineta bicolor*; Knight et al. 2018) and, notably, the congeneric blackpoll warbler (*Setophaga striata*; DeLuca et al. 2019). While blackpoll warblers are known for their incredibly long migrations, including a nonstop transoceanic flight to South America [60], myrtle warblers are typically characterized as “short-distance migrants” [61,62]. However, we found that—rather than taking the shortest route to the Pacific Coast wintering area—most myrtle warblers breeding in Anchorage traveled over 11,000 km round-trip to winter on the Gulf Coast, which is longer than the migration distances of many species considered “long-distance migrants” in other studies. Based on this result, we emphasize the importance of considering intraspecific differences in migration distance in comparative studies, and the need for more studies characterizing population-level variation in migration routes, wintering areas, and migration distance.

A number of studies have identified cases in which birds breeding in northwestern North America follow an eastern migration route, even if it seems “sub-optimal” or crosses a geographic barrier. This phenomenon has been especially well-studied in the *Catharus* thrushes, where northwestern populations of veeries (*Catharus fuscescens*) [63,64], hermit thrushes (*Catharus guttatus*) [65], and Swainson’s thrushes (*Catharus ustulatus*) [3,66,67] follow eastern migratory routes thought to follow the path of historical range expansion following the retreat of glaciers after the Last Glacial Maximum. The similarity in migration routes between these species, blackpoll warblers, and the myrtle warblers in this study suggests that the migration routes of parulid warblers breeding in Alaska may have been shaped by similar processes.

While most individuals in the present study seemed to follow this shared eastern migration route, a subset of birds may have followed an alternate migration route to the Pacific Coast. Populations in other species have been identified where a proportion of individuals migrate to a different wintering area, such as Eurasian blackcaps (*Sylvia atricapilla*) breeding in continental Europe that recently began migrating to Great Britain instead of southern Europe or Africa [4,22,68]. In tree swallows, Knight et al. [58] found that while most individuals breeding in Fairbanks, Alaska migrated east, some took a western route through Baja California to winter in Mexico. Such variation appears to be present in myrtle warblers breeding in northwestern North America, but more work is needed to determine whether a greater proportion of birds migrate to the Pacific Coast in other, unstudied breeding populations and whether Pacific Coast wintering has fitness benefits for these individuals.

### (c) Pressure geolocation improved location estimates and facilitated fine-scale characterization of migration timing and flight altitude

Methodologically, we found that while both light-level and atmospheric pressure data produced similar broad estimates of migration route, wintering area, and migration timing, pressure data allowed us to characterize migration route and timing at a much finer scale with less uncertainty, while also providing additional information about flight altitude and behavior. The spring migration routes inferred from light data using GeoLight were highly similar to those identified using pressure data in GeoPressureR for most of the geolocators. However, fall migration routes could not be inferred from light data, because fall migration overlapped the weeks around the equinox, resulting in high latitudinal error. Another benefit of using pressure data is that the changes in pressure due to migratory flights allowed us to identify every stop made by the bird, which would not be possible using light data alone. We were able to infer the locations of most of these stationary periods with relatively low uncertainty, although the geographic likelihood maps for some of the shorter stationary periods were broad, because a short pressure timeseries limits the model’s ability to constrain possible locations. The wintering areas identified using either light or pressure data alone largely overlapped, but the highest likelihood regions overlapped more closely in longitude than latitude (electronic supplementary material, figures S8 and S9, table S3), due to the higher error in latitude estimates common to light-level data [69]. The wintering areas inferred using GeoLight exhibited much higher uncertainty (i.e. a wider spread of location estimates), likely as a result of both inherent error in light data due to shading, and because the changepoint model used to identify stationary periods in some cases merged short stops at the end of fall migration and beginning of spring migration with the wintering period.

In addition to improving location and timing estimates, the inclusion of pressure data also provided insights into aspects of myrtle warbler behavior that could not be studied using traditional geolocators collecting light data alone. From pressure readings taken during migratory flights, we were able to estimate that the birds flew at a median altitude of ∼840m above sea level, reaching a maximum altitude of 2,900m a.s.l., and we were able to document changes in altitude over the course of each flight. Most estimates of migratory flight altitude have been obtained for species breeding in Europe, many of which have been found to fly at higher altitudes than the myrtle warblers in our study [70–74]. The most comparable of these species to the myrtle warbler, the great reed warbler (*Acrocephalus arundinaceus*), flew at a median flight altitude of 1,150–1,630m a.s.l. and reached a maximum altitude of ∼6,500m a.s.l. [71,74,75]. Few studies of migratory flight altitude have been conducted in North America, and most have used radar to broadly characterize flight altitude for the aggregate of birds migrating over a region. Radar [76,77] and radio-telemetry studies [78], along with our pressure data, suggest that bird migration in North America may occur at a lower altitude than in Europe, potentially due to differences in topography, geographic barriers, or wind patterns.

Pressure data can also be used to detect fine-scale altitudinal movements throughout the annual cycle. In our study, we observed cyclical pressure changes in some individuals during winter, with one individual spending nighttime periods at an elevation 50m higher than the daytime elevation. This individual may have been traveling to a higher-elevation nocturnal roost site and foraging at a different location during the day, a behavior that has been recorded in myrtle warblers once previously [79]. Because pressure geolocators can provide fine-scale altitude data for individual birds, this method is highly valuable for characterizing elevational movements and flight altitude in different species and for investigating what factors, such as weather, topography, or urbanization, may affect flight altitude throughout the annual cycle.

## CONCLUSIONS AND FUTURE DIRECTIONS

We found that, contrary to expectations, most myrtle warblers breeding in Alaska and northern British Columbia migrated over 11,000 km round-trip to winter in the southeastern United States, rather than taking a shorter, alternate migration route to the Pacific Coast wintering area. The route followed by the geolocator-tracked birds was similar to that taken by other songbirds breeding in northwestern North America and is consistent with the previously proposed hypothesis that current migration routes reflect range expansions after the Last Glacial Maximum. Stable isotope results suggested that a small subset of birds breeding in both Alaska and British Columbia may winter on the Pacific Coast, with no migratory divide present but the proportion of Pacific-wintering birds increasing farther west.

Future work should focus geolocator deployment or stable isotope sample collection in unsampled regions of the northwestern breeding range or in the Pacific Coast wintering area to determine where more Pacific-wintering birds breed. Future studies could also investigate whether there is a genetic basis to the differences in migration route between myrtle warblers wintering in the southeast versus the Pacific Coast, and whether birds that winter on the Pacific Coast practice assortative mating on the breeding grounds, as has been observed in other birds that migrate to an alternate wintering area [22].

Methodologically, we found that the use of multi-sensor geolocators that record atmospheric pressure data allowed us to characterize migration routes and timing at a much finer scale than using traditional light-level data alone, while also providing information on flight altitude and fine-scale elevational changes during the wintering period. We recommend that researchers planning geolocation studies employ tags with barometers, as the additional pressure data greatly improved our ability to characterize migration and behavior over the annual cycle for only a small increase in the cost and weight of the tag. Fine-scale tracking of individual birds from a greater diversity of species will reveal patterns of intraspecific variation in migratory behavior and help to increase understanding of relationship between migration and geographic range.

### Ethics

Samples for this study were collected under protocols approved by IACUC at Penn State University (201900879) and UC Riverside (20210012) and under USGS federal bird banding permit #24042, Environment and Climate Change Canada federal bird banding permit #10897C, Alaska State Scientific Permit #22-130, British Columbia Ministry of Forests Wildlife Act Permit #FJ22-624396, and Alberta Ministry of Environment and Protected Areas Permit #21-253.

### Data accessibility

The geolocator data generated in this study, including raw light and pressure data and location estimates, are available on Movebank (ID #4549104814). The scripts used to process the geolocator data are available on Github (https://github.com/sszarmach/myrtle_geo_pressure). Sampling information and morphological data are provided as electronic supplementary material (Supporting Datasets 1 and 2).

### Authors’ contributions

SJS, DPLT, AB, and BMV contributed to the study conception and design. SJS, DPLT, and AB acquired funding for the project. SJS, JKB, and MM conducted the field work and sample collection. SJS performed the stable isotope sample preparation and the geolocator and stable isotope data analysis. The first draft of the manuscript was written by SJS, and all authors reviewed the manuscript, contributed to subsequent drafts, and approved the final draft.

### Funding

This research was funded by student research grants awarded to S.J.S. from the American Ornithological Society, the Animal Behavior Society, and the Wilson Ornithological Society (Paul A. Stewart Grant). This work was also funded by a Pennsylvania State University Science Achievement Graduate Fellowship awarded to S.J.S., the Department of Biology and Huck Institute of Life Sciences at Pennsylvania State University, and the University of California Riverside Department of Evolution, Ecology, and Organismal Biology. Collection of samples used in the stable isotope analysis was funded by an Alberta Conservation Association Grant in Biodiversity awarded to Daniel Pierce.

## Supporting information

Supporting Dataset 1 - Geolocator and Control Bird Sampling

Supporting Dataset 2 - Isotope Sampling

## Acknowledgements

We thank Daniel Pierce, Alana Demko, Olga Lansdorp, German Lagunas-Robles, Michael Moretti, Marie Palanchon, and Allison Patterson for the collection of feather samples used in the stable isotope analysis. We are grateful to the Cornell Stable Isotope Laboratory, in particular to Kim Sparks, who provided training in stable isotope sample preparation. We also thank Henry Streby for providing training in geolocator harness construction and geolocator deployment. We are grateful to Raphael Nussbaumer for assistance with the GeoPressureR package. We also thank Andrew Wood for support with project logistics. Special thanks to Emily Griffith for creating the myrtle warbler illustrations used in Figures 1 and 2.

