## Supplementary Figures and Tables for "Migration strategies of a high-latitude breeding songbird (*Setophaga coronata coronata*) revealed using multi-sensor geolocators and stable isotopes"

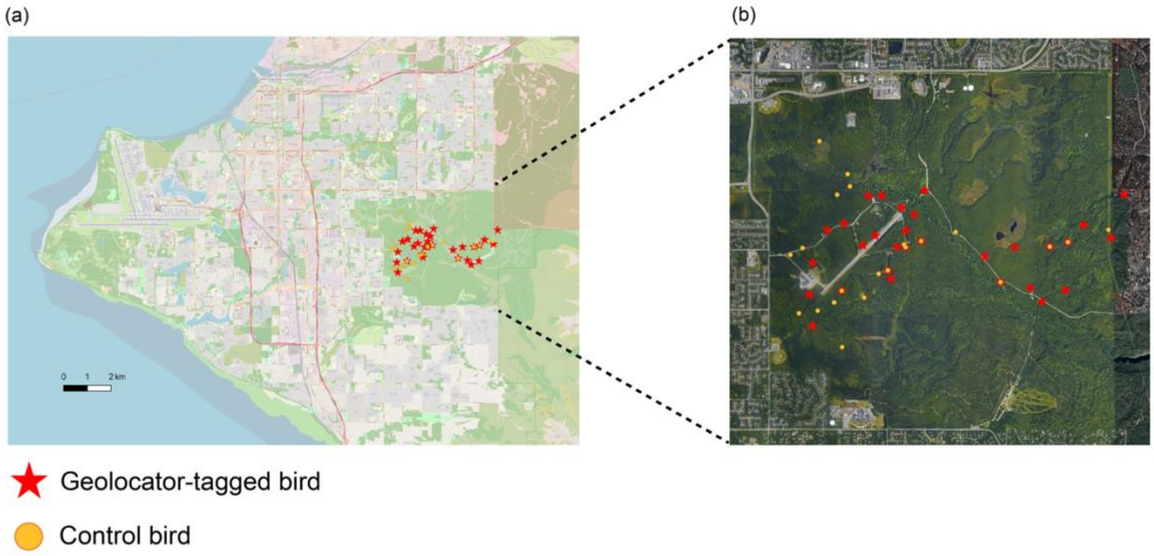


**Figure S1.** Sampling area for geolocator study. (a) Location of study area within Anchorage, AK. (b) Locations where myrtle warblers were captured within Far North Bicentennial Park. Red stars indicate sites where myrtle warblers were tagged with geolocators (*n* = 30), yellow circles represent sites where “control” birds received a color band, but not a geolocator (*n* = 25). Base map: Open Street Map Standard (obtained through QuickMapServices QGIS plugin), retrieved 22 September 2022.

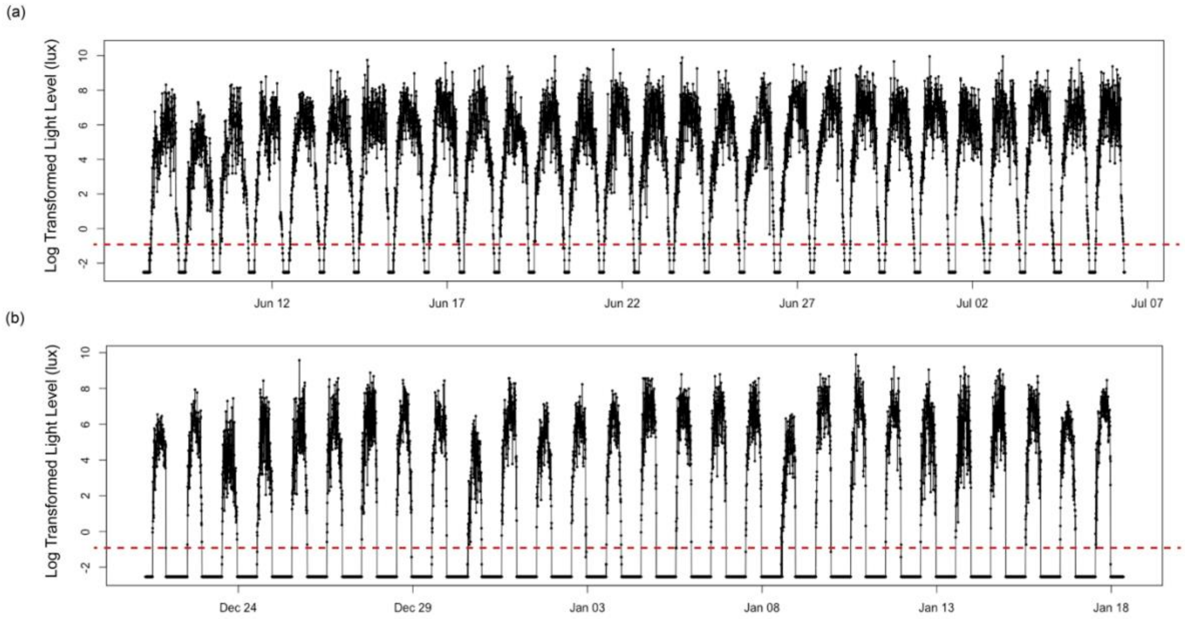


**Figure S2.** Light level data from one geolocator over two time periods: (a) summer on the breeding ground (9 June 2022—6 July 2022), and (b) winter on the non-breeding ground (22 December 2023—18 January 2023). The twilight threshold used for GeoLight analyses is shown by the dashed red line. Light data is log transformed to improve visualization.

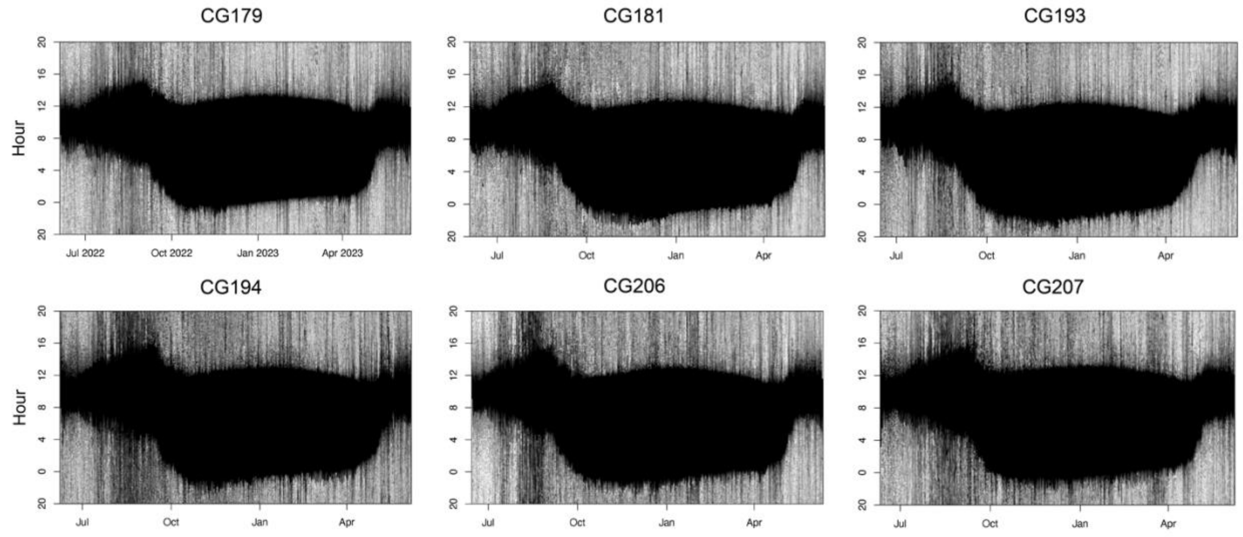


**Figure S3.** Light level measured over the full year for each of six geolocators. Each vertical line represents light levels over one day, with hours of the day represented along the y-axis. The nighttime period has been centered in each plot. Dark pixels represent complete darkness, while increasingly white pixels represent higher levels of light. Sudden shifts in the timing of darkness (night) are indicative of movement to a different location.

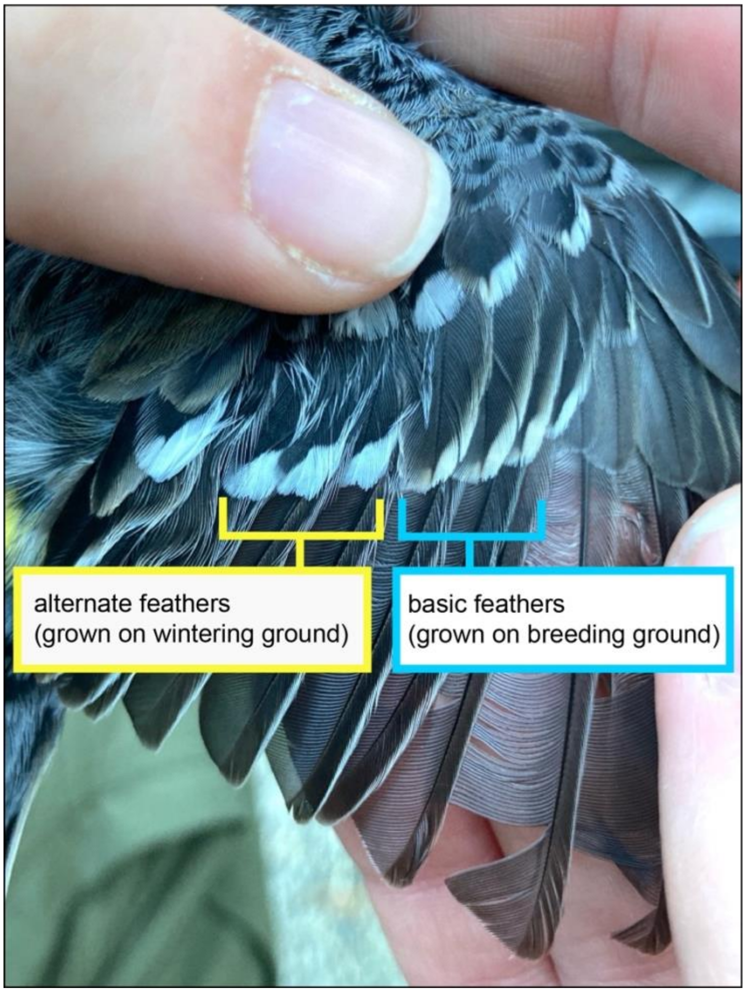


**Figure S4.** Photograph of myrtle warbler wing exhibiting clear molt limit between alternate greater covert feathers, likely grown on the wintering ground, and basic feathers, likely grown on the previous year's breeding ground. Alternate feathers, grown more recently, are darker black with brighter and more extensive white tips.

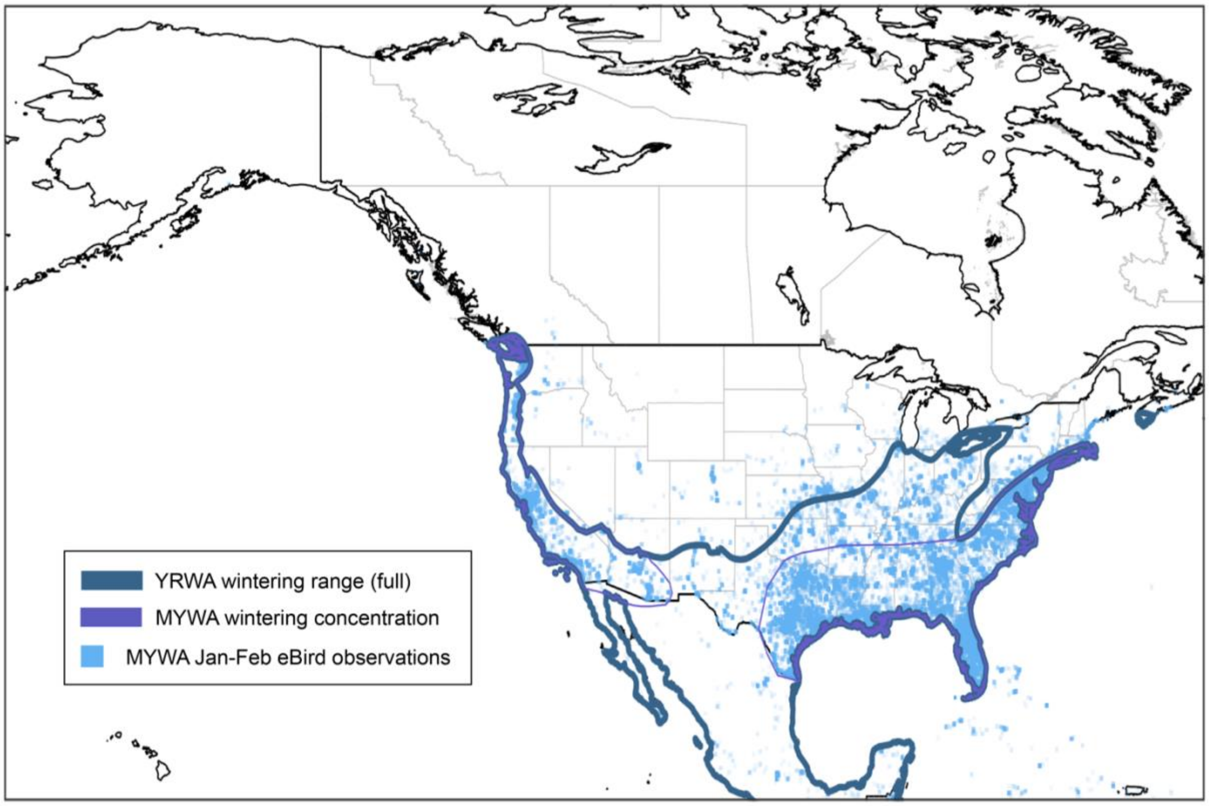


**Figure S5.** Map of myrtle warbler wintering areas. The dark blue outline represents the full myrtle warbler wintering range, modified from BirdLife International yellow-rumped warbler range shapefiles. The purple outlines surround the regions of greatest myrtle warbler wintering abundance, based on eBird observations (blue points) and the eBird Status and Trends abundance map. These wintering concentration polygons were used for assessing differences in precipitation stable hydrogen ratios ($\partial^2$H) between the two wintering areas, and for calculating odds ratios of feather origins from the East Coast versus West Coast.

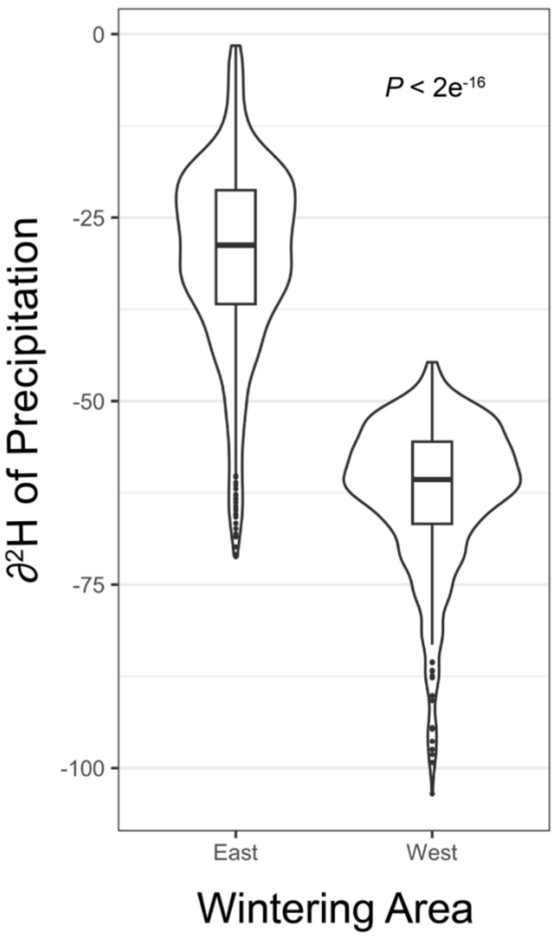


**Figure S6.** Precipitation stable hydrogen isotope values ($\partial^2H$) sampled from 1000 random points in the myrtle warbler eastern wintering range vs. 1000 points in the western wintering range. The western wintering range exhibited significantly lower $\partial^2H$ values ($P < 2e^{-16}$), so myrtle warbler feathers grown on the Pacific Coast wintering ground are expected to have lower stable hydrogen ratios compared to those grown on the Gulf and Atlantic Coasts.

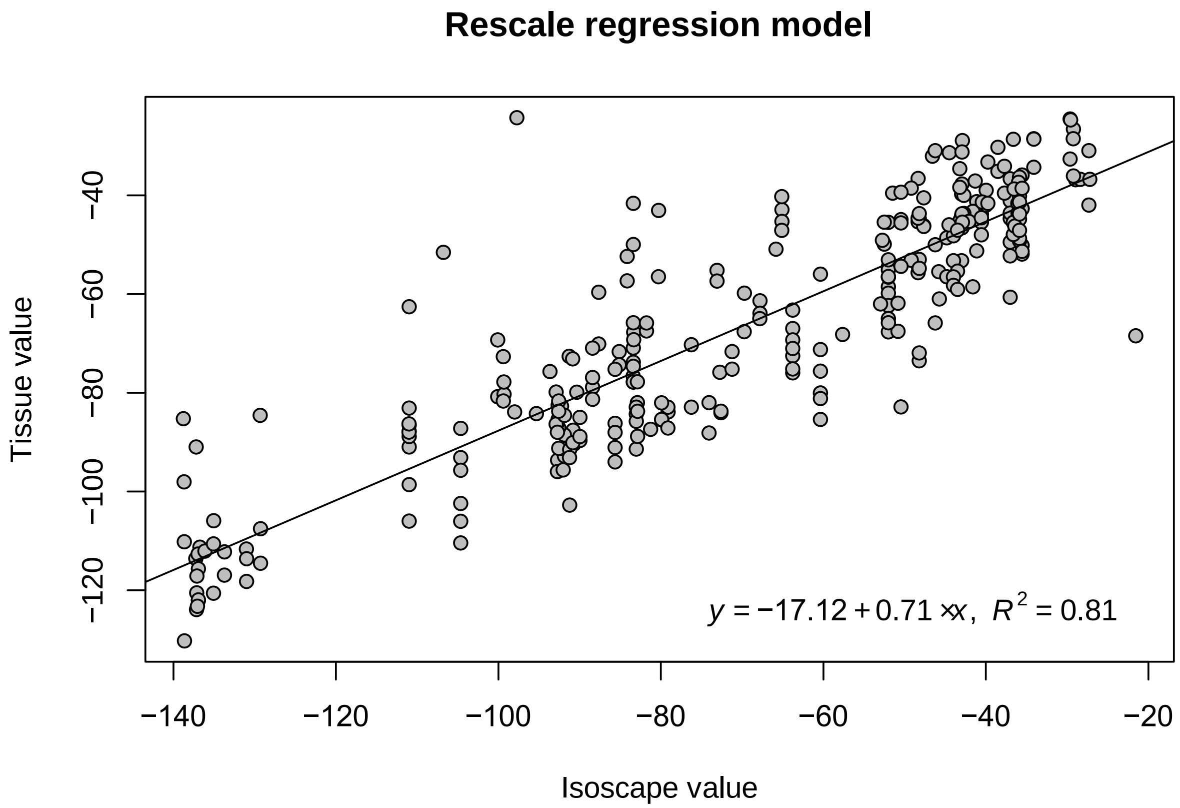


**Figure S7.** Transfer function between precipitation stable isotope values ($\partial^2H_p$) and feather isotope values ($\partial^2H_f$) generated from 313 feather samples from 19 parulid warbler species with known origins.

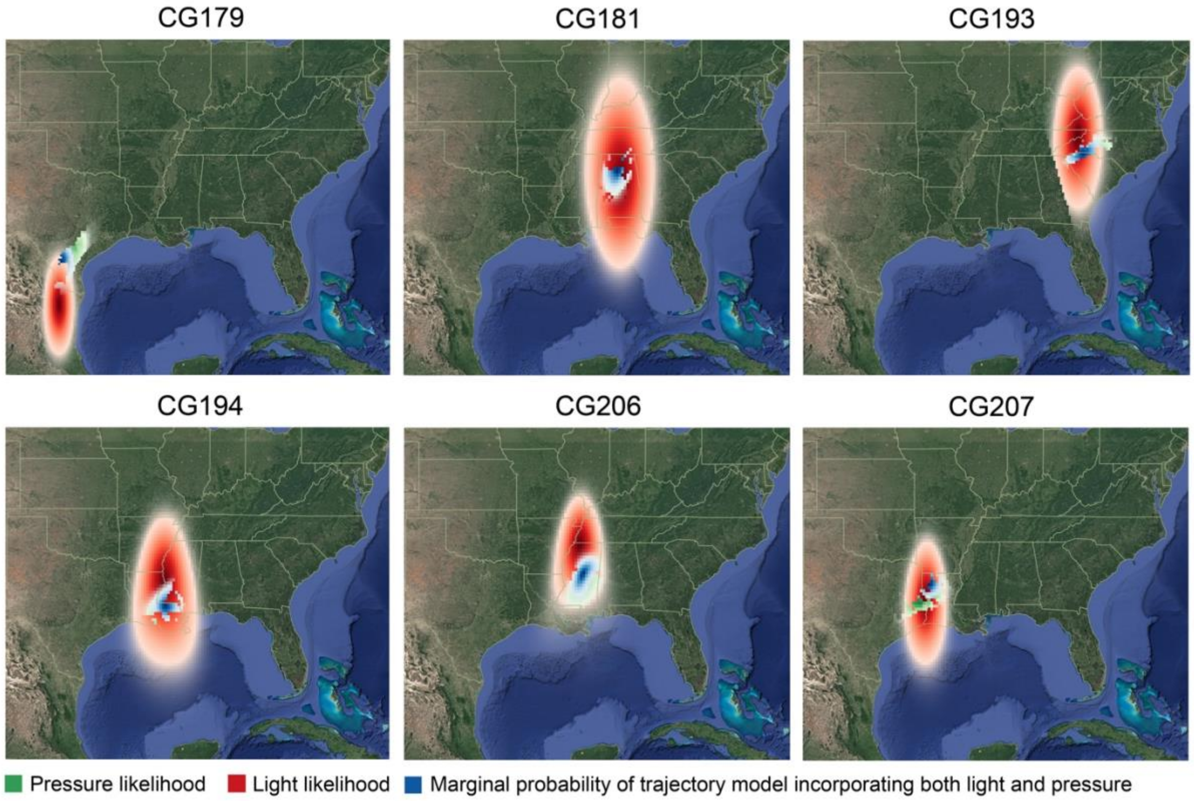


**Figure S8.** Likelihood maps produced using GeoPressureR for the longest winter stationary period of each bird. Red ovals represent likely wintering areas based on light data alone, green pixels represent likely wintering areas based on atmospheric pressure data alone, and blue pixels show the marginal probability of the model incorporating both light and pressure data. Darker colors represent higher likelihoods for all three products. For some tags (e.g. CG181, CG193, CG194) the pressure likelihood and marginal probability of the full model show high overlap, so few green pixels are visible. Base map: Google Satellite Hybrid (obtained through QuickMapServices QGIS plugin), retrieved 2 April 2024.

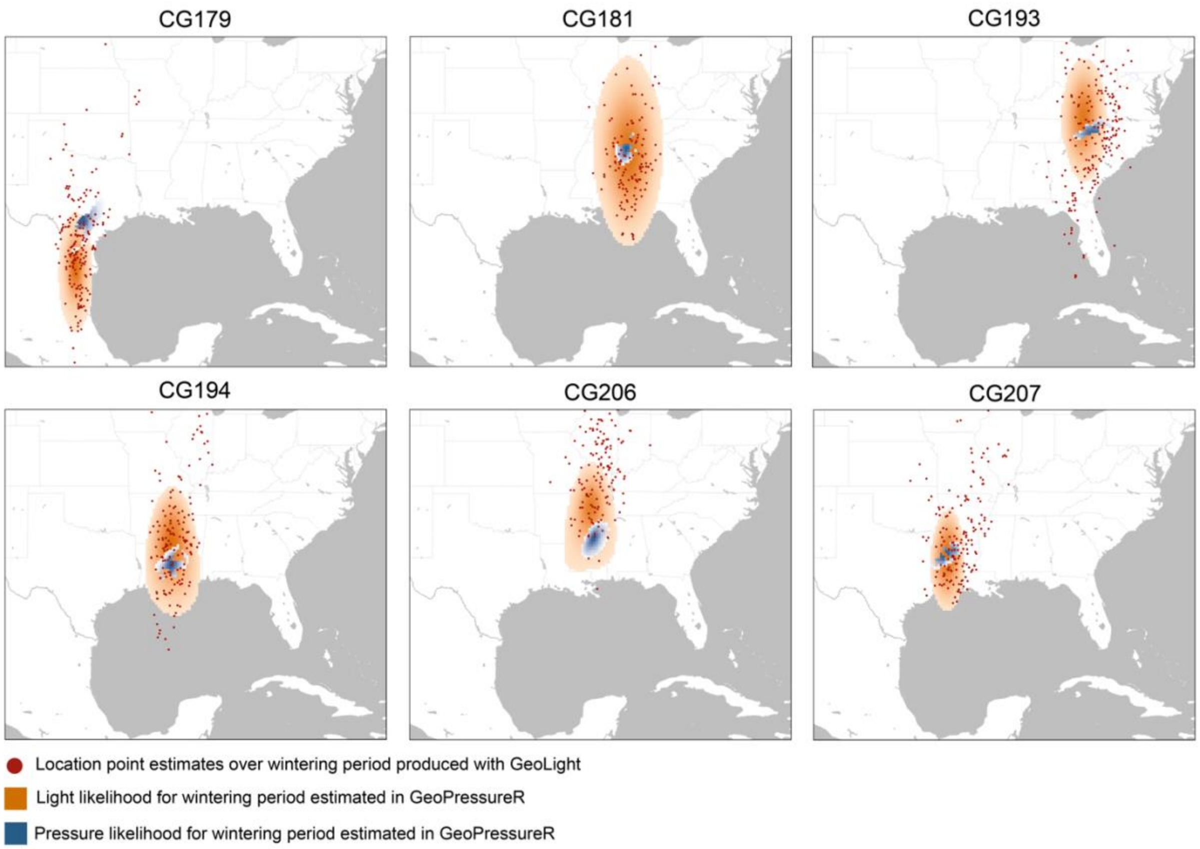


**Figure S9.** Comparison of wintering areas for six myrtle warblers estimated using three methods: light level data alone analyzed using the threshold method in GeoLight (red), light level data using GeoPressureR (orange), and pressure data alone using GeoPressureR (blue). Darker colors represent higher likelihoods for all three products. Base map: Stamen Design Toner Background, data from OpenStreetMap, retrieved 3 April 2024.

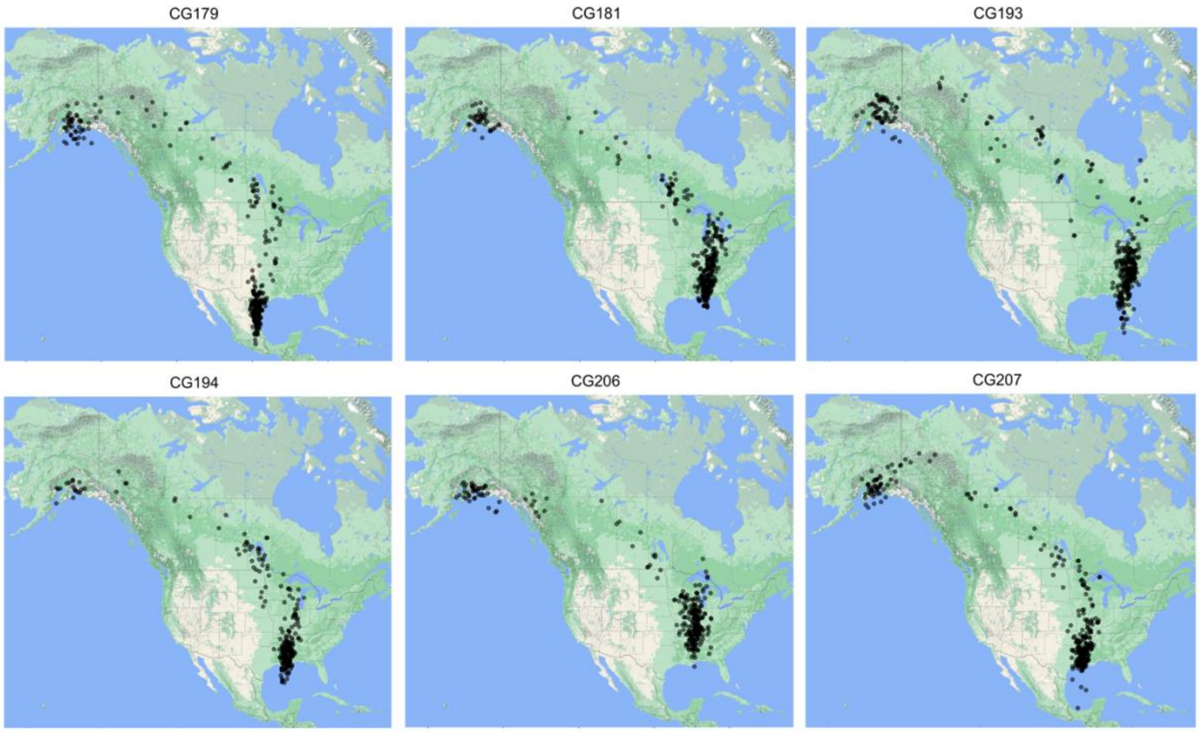


**Figure S10.** Location point estimates for the breeding ground, spring migration path, and wintering ground inferred from geolocator light-level data using the threshold method in the GeoLight package. Each panel shows the track of one bird. The fall migration path was not plotted due to high error in latitude estimates resulting from equal day lengths near the autumn equinox, and points from within three weeks before and after the spring equinox were also removed for the same reason. Map data ©2024 Google, INEGI.

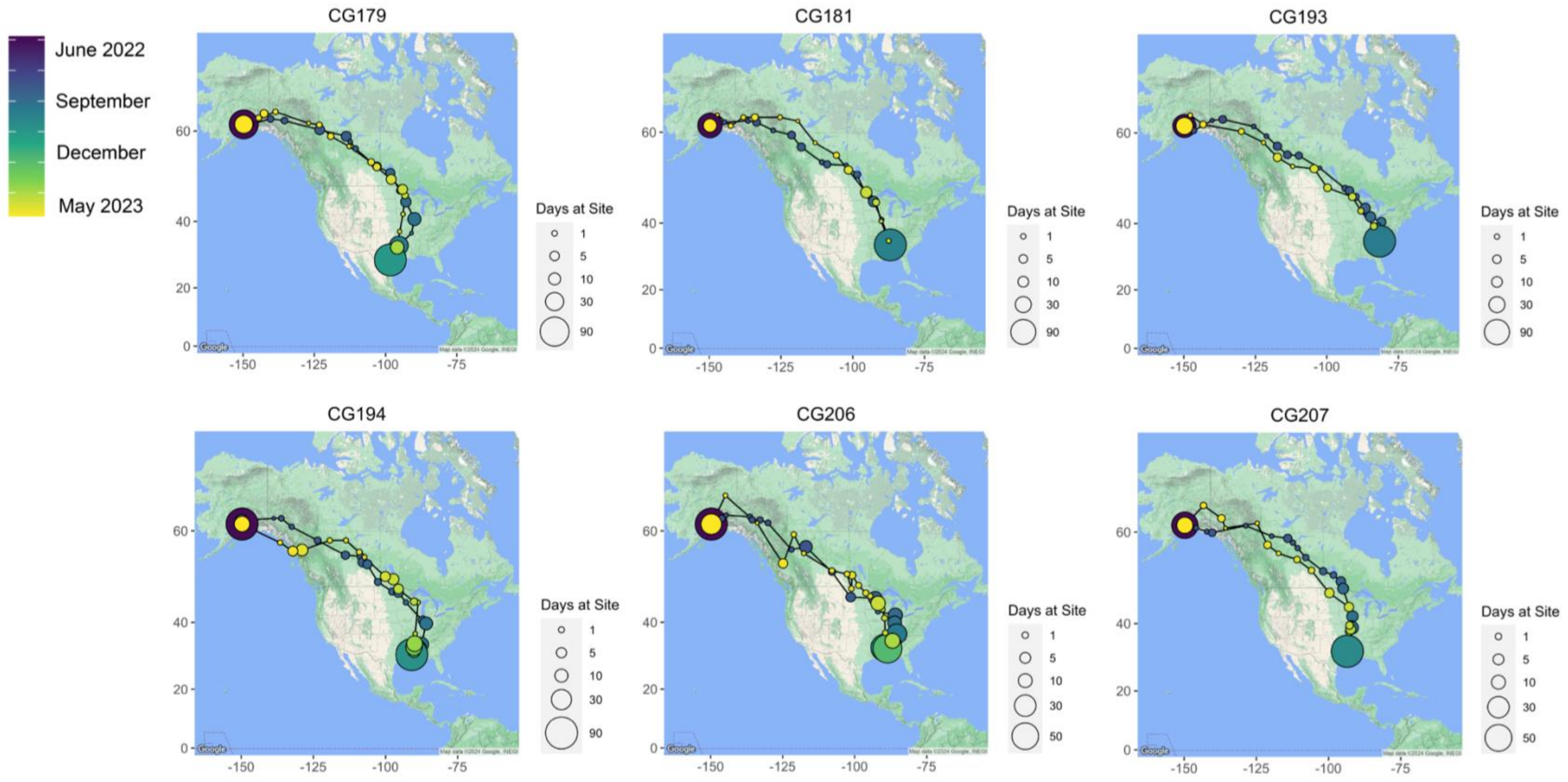


**Figure S11.** Year-long migration routes for six myrtle warblers inferred from pressure data alone using the GeoPressureR package. The size of each circle represents the amount of time spent at a site, and the color represents time of year (fall migration: blue, spring migration: yellow).

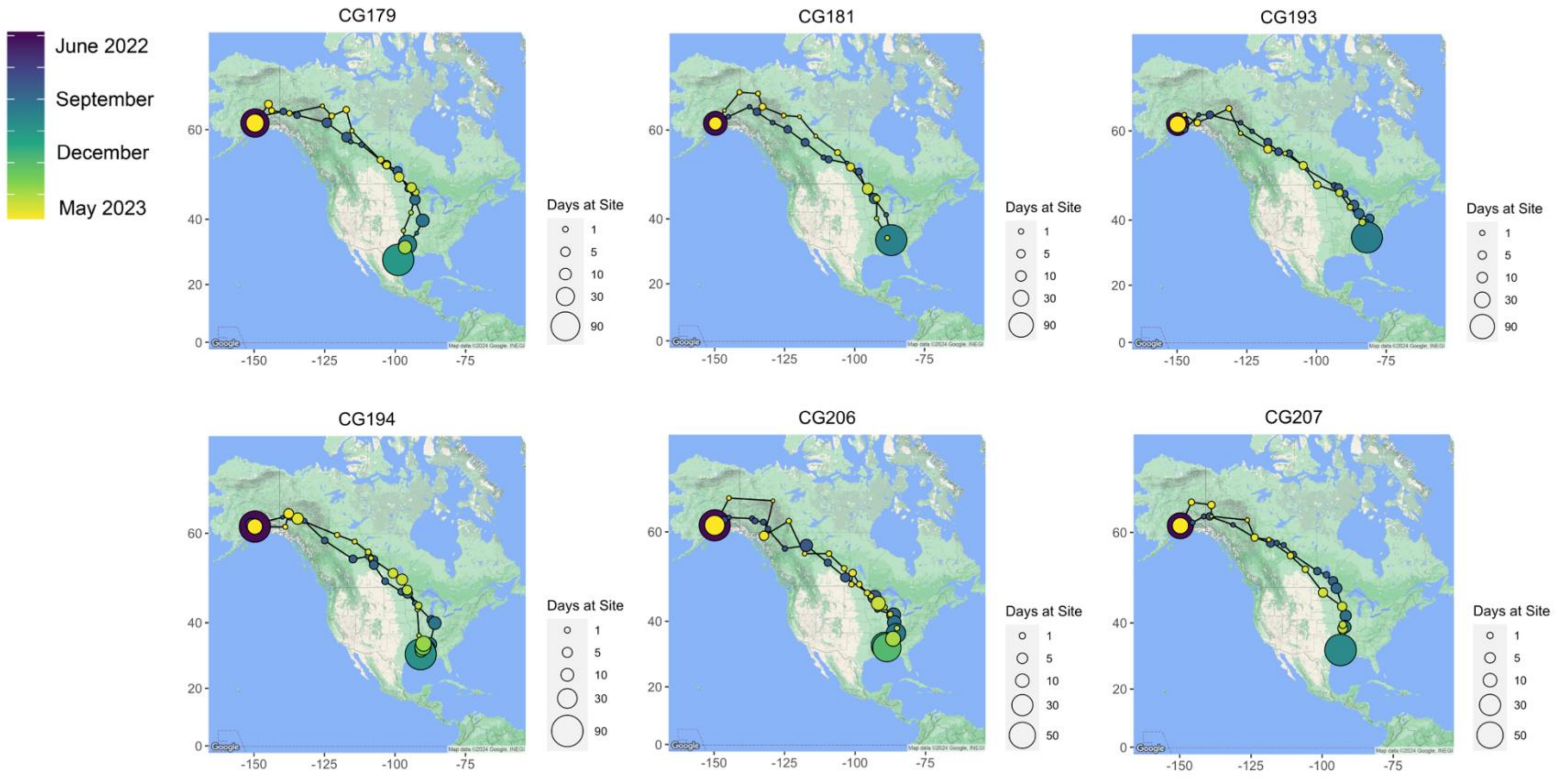


**Figure S12.** Year-long migration routes for six myrtle warblers inferred using light-level data, pressure data, and a movement model using the GeoPressureR package. The size of each circle represents the amount of time spent at a site, and the color represents time of year (fall migration: blue, spring migration: yellow).

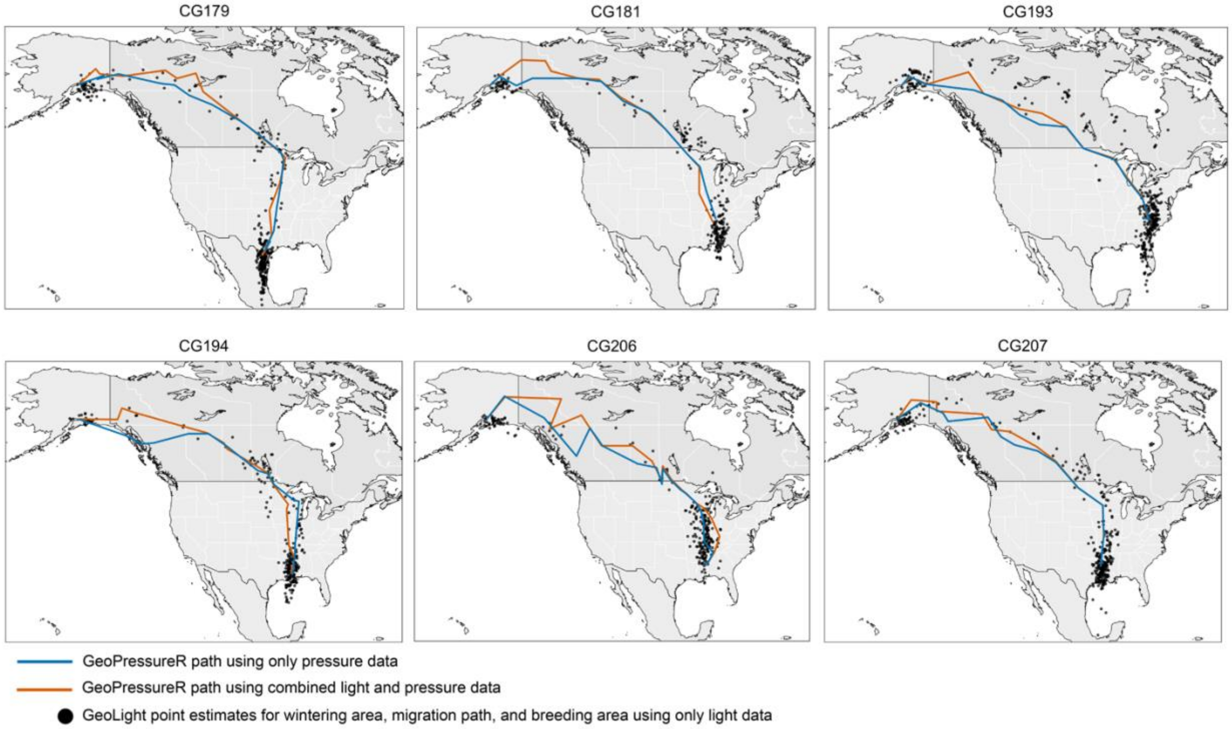


**Figure S13.** Comparison of spring migration paths for six geolocator-tracked myrtle warblers estimated using three methods: light level data alone analyzed using the threshold method in GeoLight (black points), pressure data alone using GeoPressureR (blue path), and a model incorporating light, pressure, and movement speed probability in GeoPressureR (orange path).

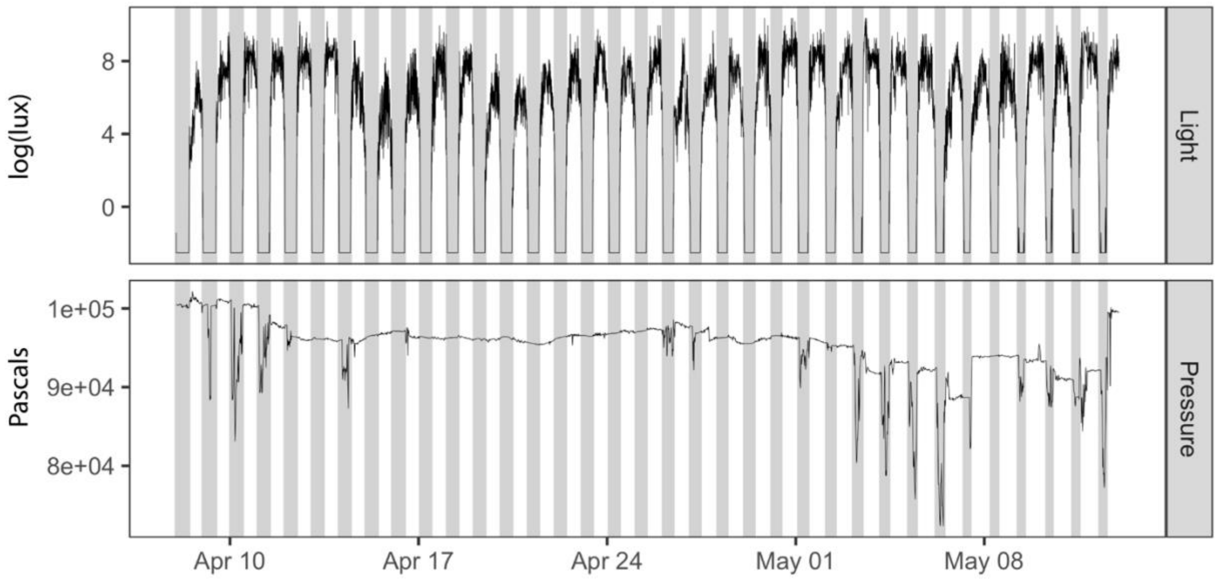


**Figure S14.** Nocturnal flight behavior of one geolocator-tracked myrtle warbler (CG181) over the spring migration period. Top panel shows light level measured by the geolocator (log transformed for better visualization). Bottom panel shows atmospheric pressure in Pascals for the same time period. Periods where the geolocator measured complete darkness are shaded grey. Large, sharp drops in pressure indicative of migratory flights almost always occur during periods of darkness, demonstrating the nocturnal migratory behavior of myrtle warblers.

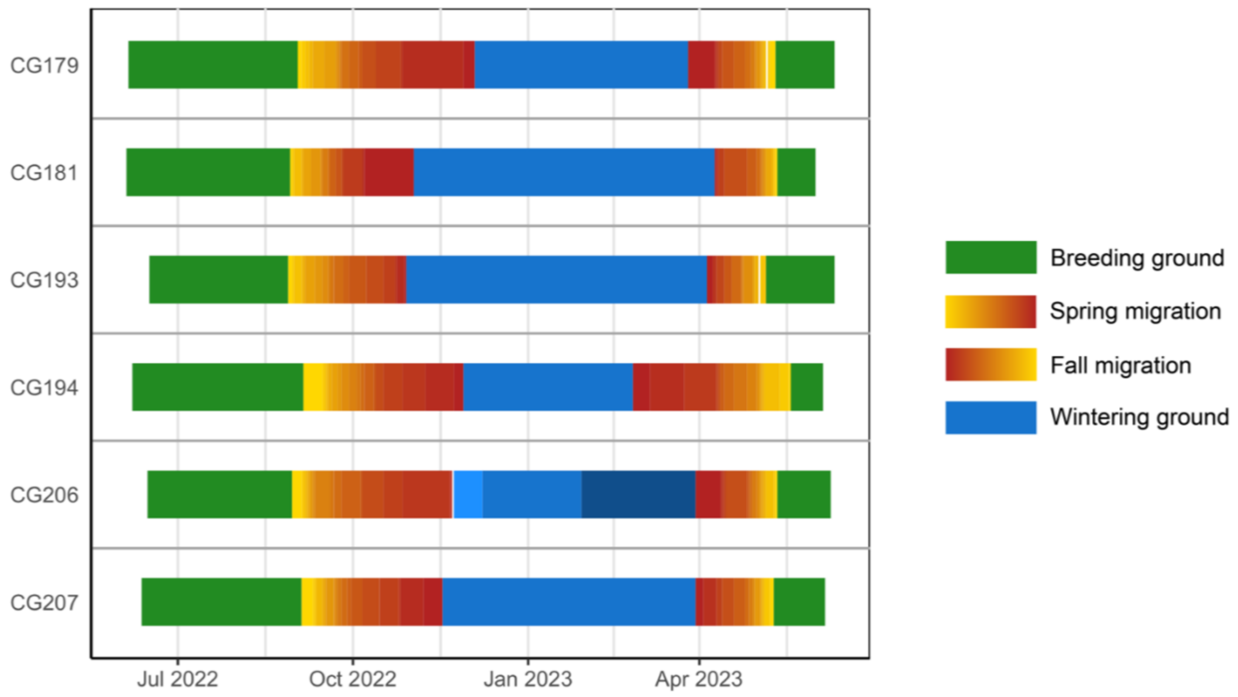


**Figure S15.** Migration timelines for six geolocator-tracked myrtle warblers estimated using atmospheric pressure data. Each horizontal bar represents the timeline for one bird. Color coding indicates time spent on the breeding ground (green), migrating (yellow/red gradient), and on the wintering ground (blue). Bird CG206 moved between three locations within 150km during the wintering period, and time spent at each of these sites is indicated using different shades of blue.

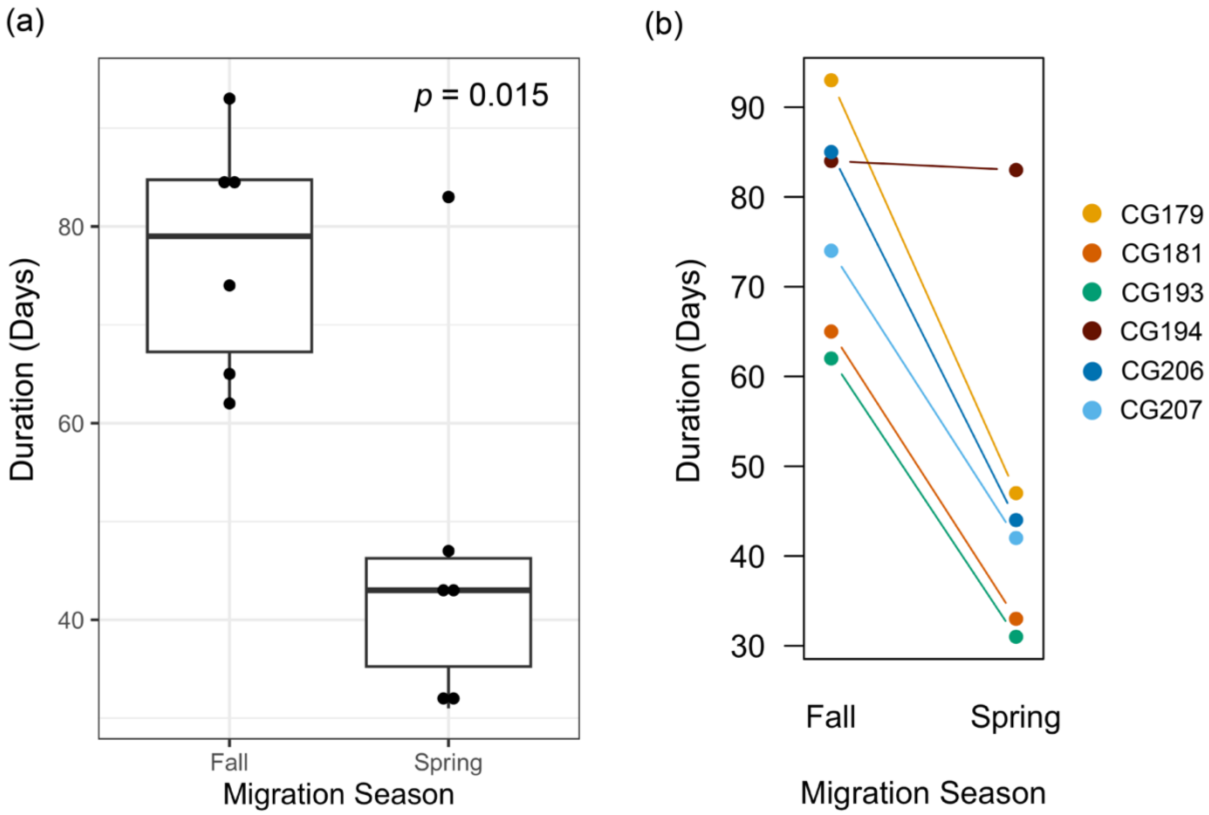


**Figure S16.** Comparison of duration of fall and spring migration in geolocator-tracked myrtle warblers. (a) Boxplots showing median and interquartile range of migration duration in fall and spring. Migration duration was significantly lower in spring ($W$=33, $p$=0.015). (b) Interaction plot showing differences in migration duration between seasons for individual birds.

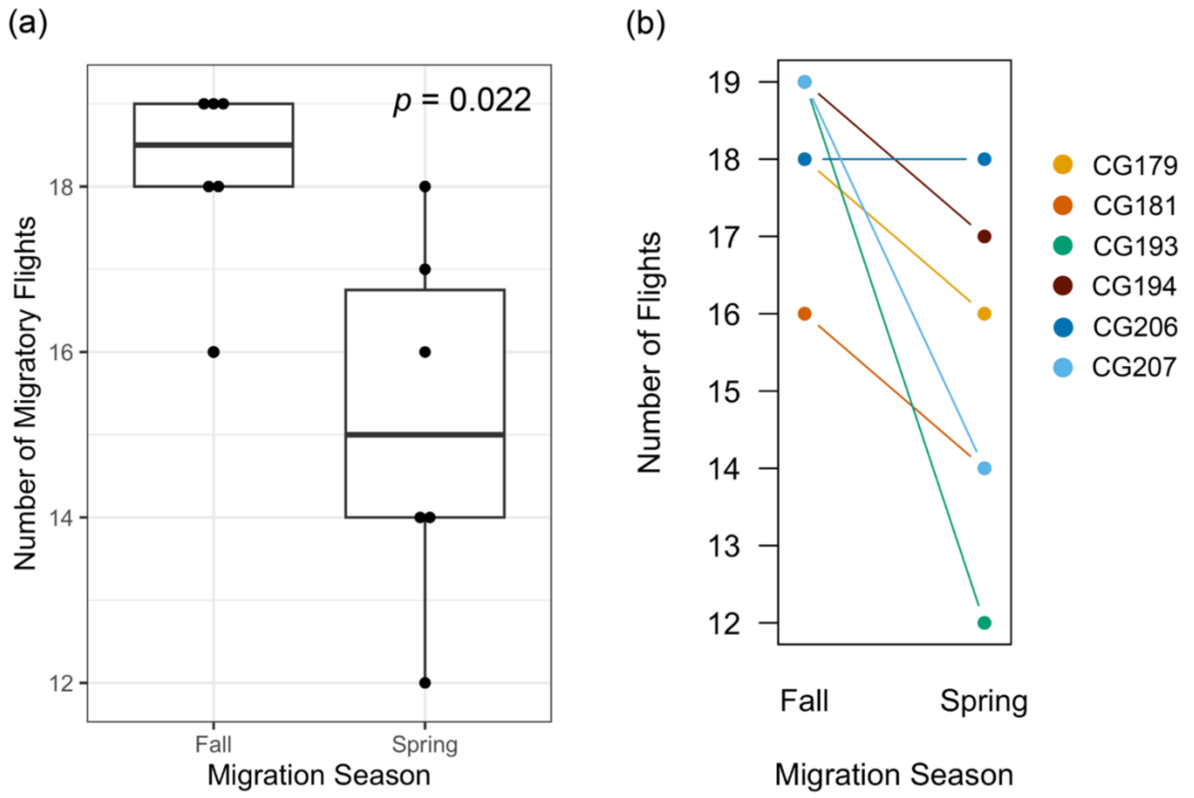


**Figure S17.** Comparison of number of flights in fall and spring migration for geolocator-tracked myrtle warblers. (a) Boxplots showing median and interquartile range of number of migratory flights in fall and spring. The number of flights was significantly lower in spring ($W$=32.5, $p$=0.022). (b) Interaction plot showing differences in the number of migratory flights between seasons for individual birds.

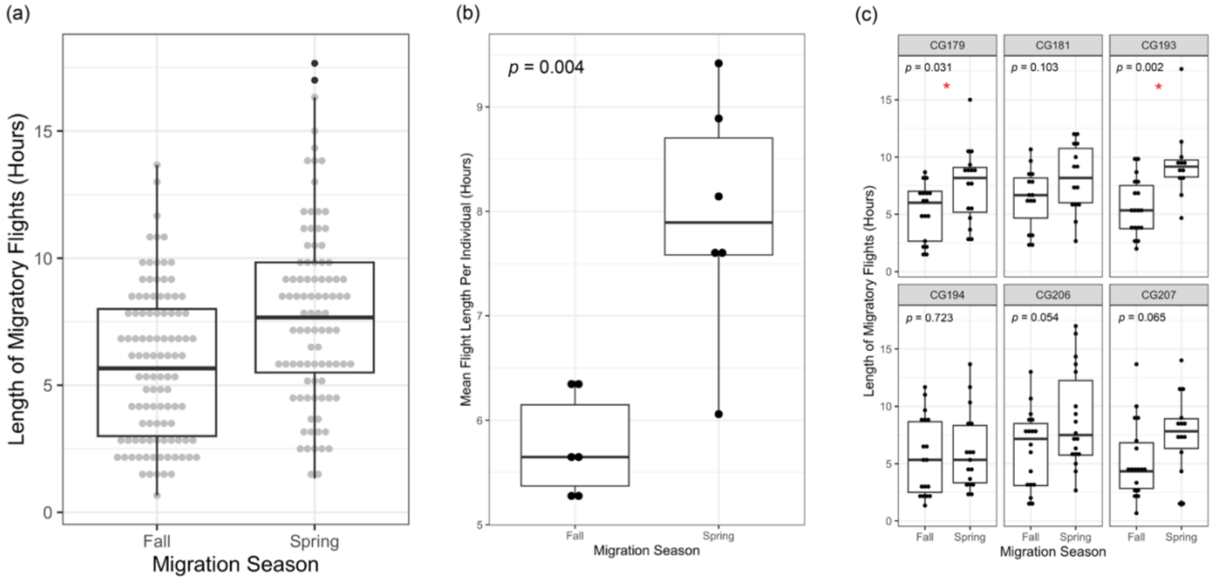


**Figure S18.** Comparison of length of migratory flights in fall and spring for geolocator-tracked myrtle warblers. (a) Boxplots showing length of migratory flights in fall vs. spring including all flights from all six birds. (b) Boxplots showing the mean flight length over a season for each bird. The mean flight length per bird was significantly higher in spring than fall ($t$ = -4.23, df = 6.7, $p$ = 0.004). (c) Boxplots showing differences in flight length between spring and fall for each individual bird.

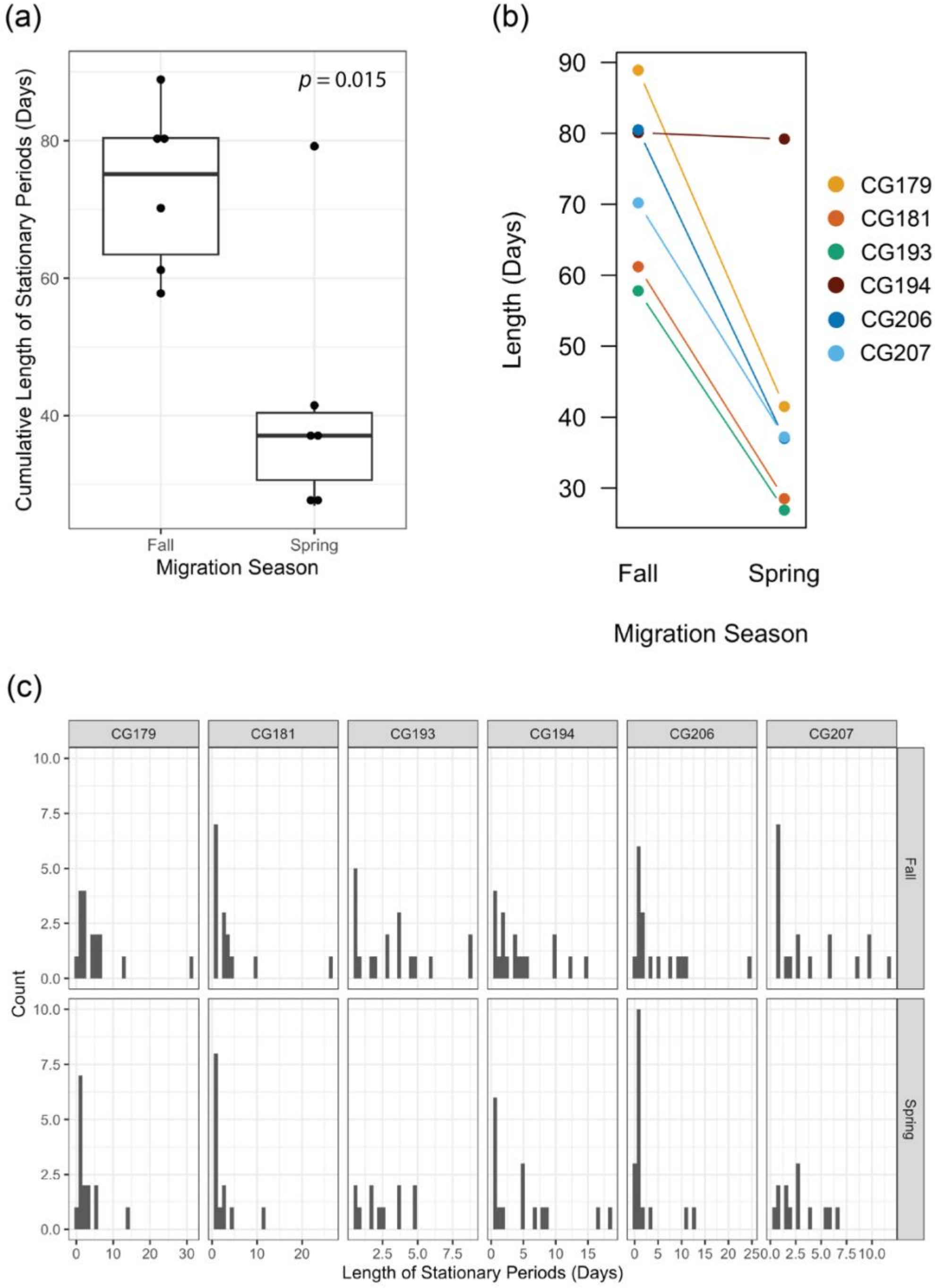


**Figure S19.** Comparison of length of stationary periods between migratory flights in fall versus spring for geolocator-tracked myrtle warblers. (a) Boxplots showing total stationary time during fall vs. spring migration (sum of lengths of all stationary periods during the season). Birds were stationary for significantly more time during fall than spring migration ($W = 33$, $p = 0.015$). (b) Interaction plot showing total stationary time in fall vs. spring migration for each individual bird. (c) Histograms of stationary period lengths for each bird in fall and spring migration.

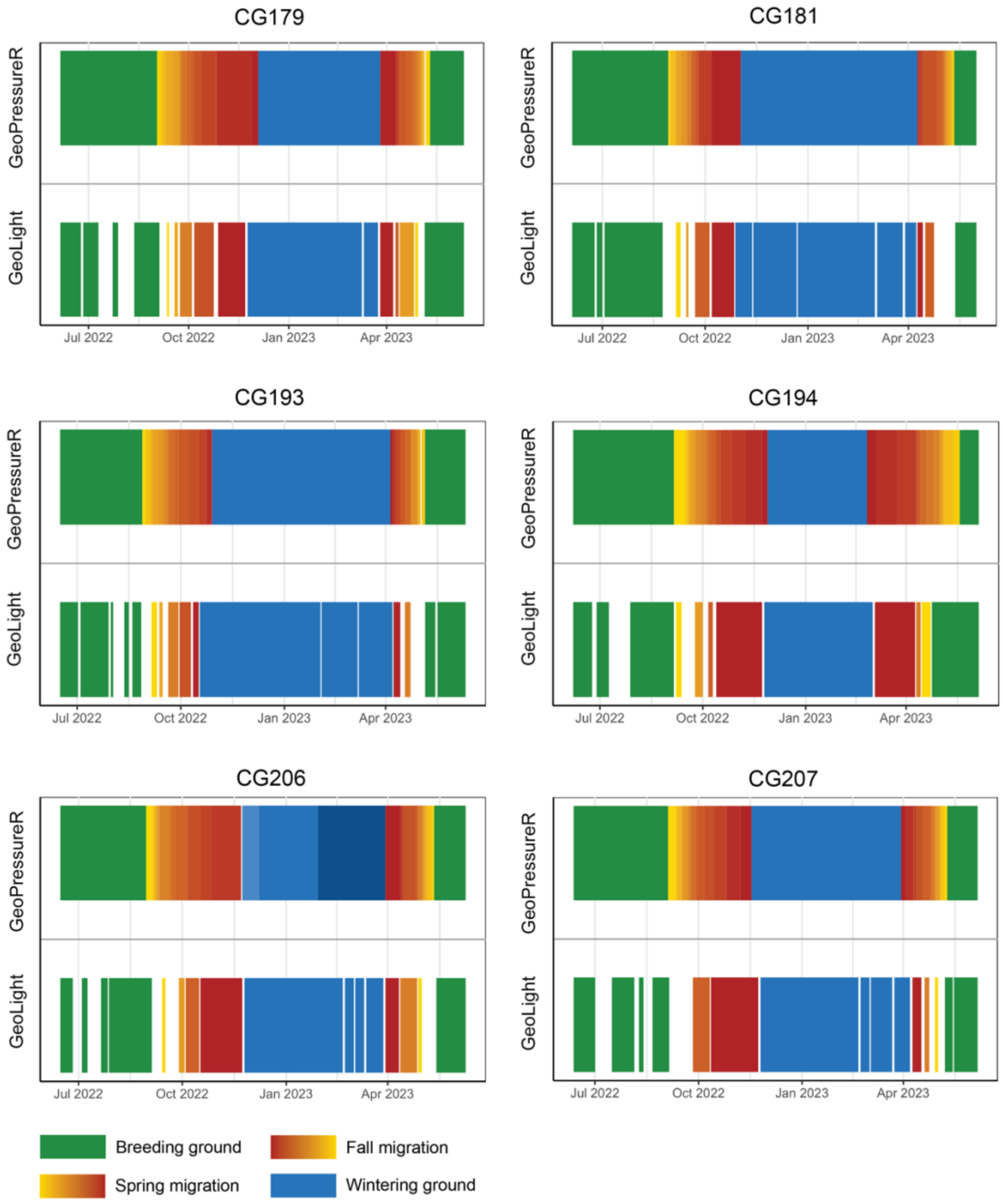


**Figure S20.** Comparisons of migration timelines generated for six geolocator-tracked myrtle warblers using atmospheric pressure data in the GeoPressureR package versus light-level data in the GeoLight R package. Each panel of two timelines represents data from one bird. Color coding indicates time spent on the breeding ground (green), migrating (yellow/red gradient), and on the wintering ground (blue). Stationary periods estimated using GeoLight that aligned with the breeding or wintering periods identified by GeoPressureR were also colored green or blue, respectively. White spaces in the GeoLight timelines are periods when the bird was not inferred to be stationary. Bird CG206 moved between three locations within 150km during the wintering period, and time spent at each of these sites is indicated using different shades of blue.

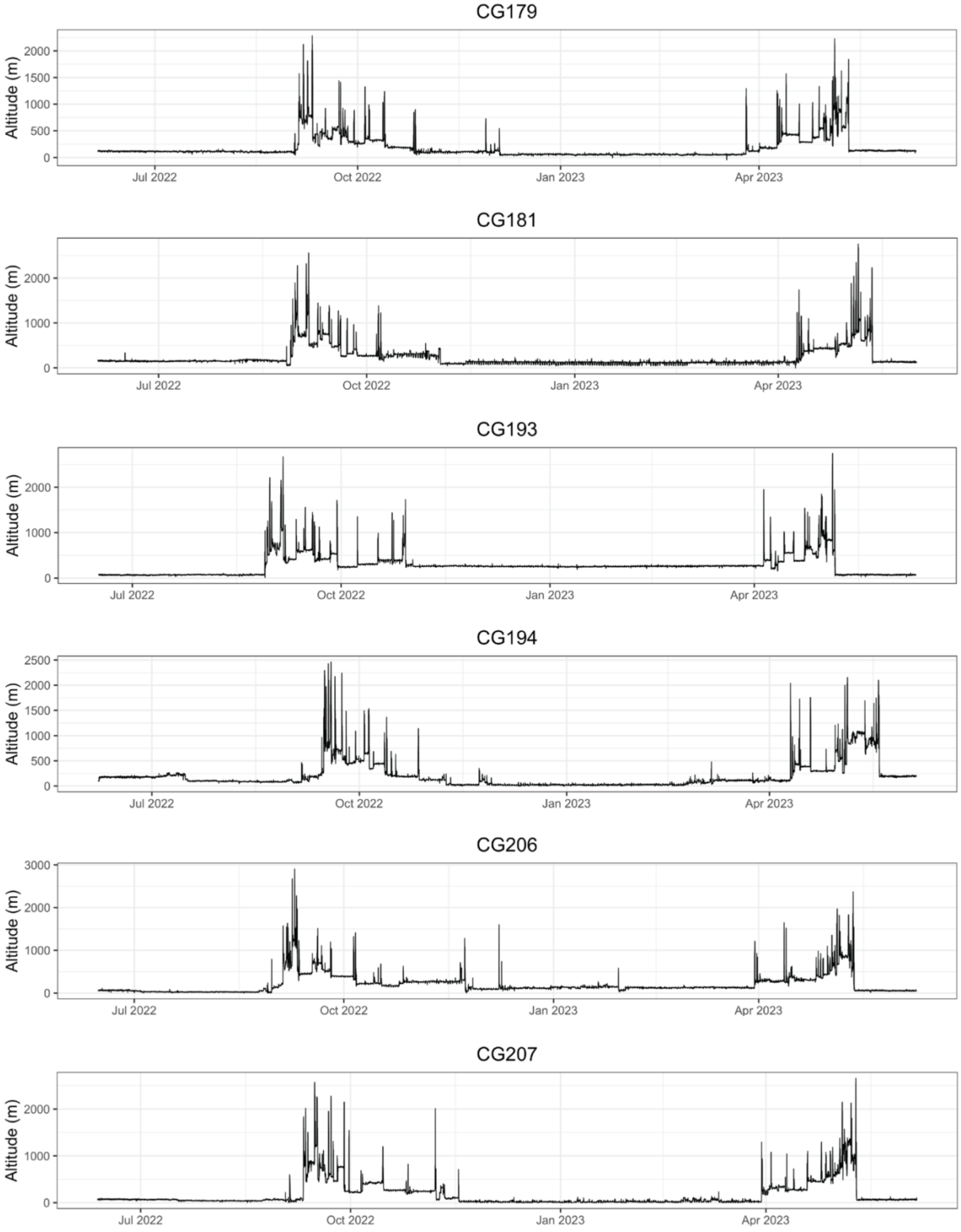


**Figure S21.** Altitude over the full year for six myrtle warblers determined from atmospheric pressure data recorded by geolocators.

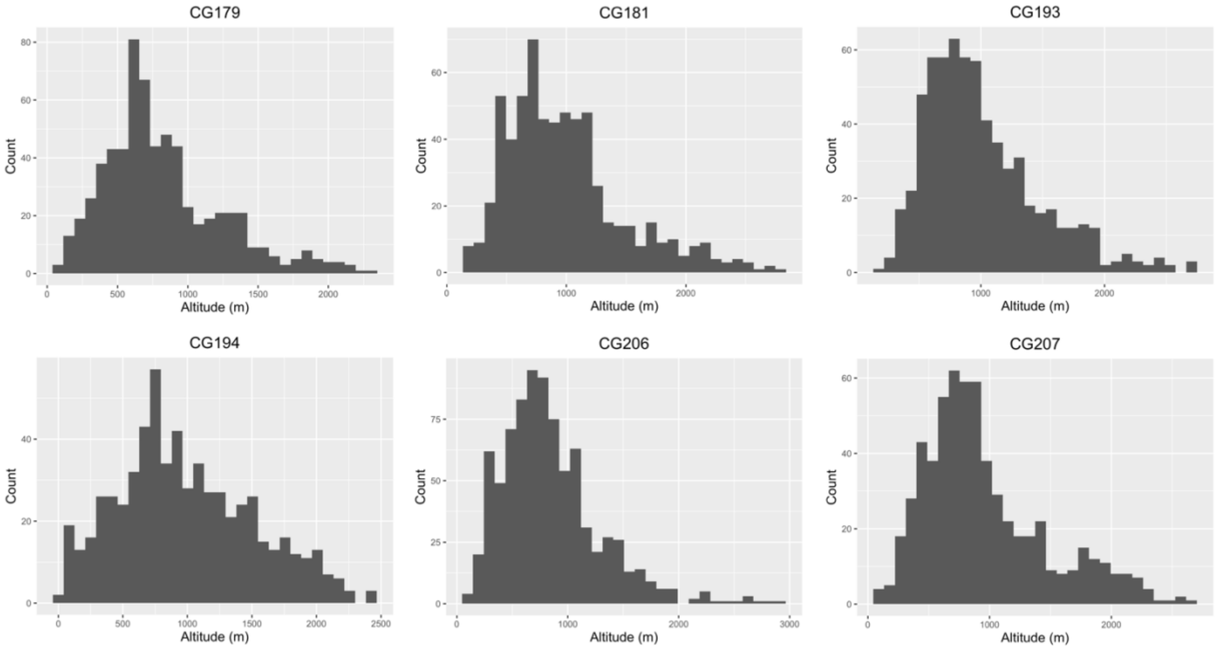


**Figure S22.** Histograms depicting distributions of flight altitude for six myrtle warblers determined from atmospheric pressure data collected by geolocators. Pressure readings were recorded every 20 minutes to characterize changes in altitude over the course of the flight.

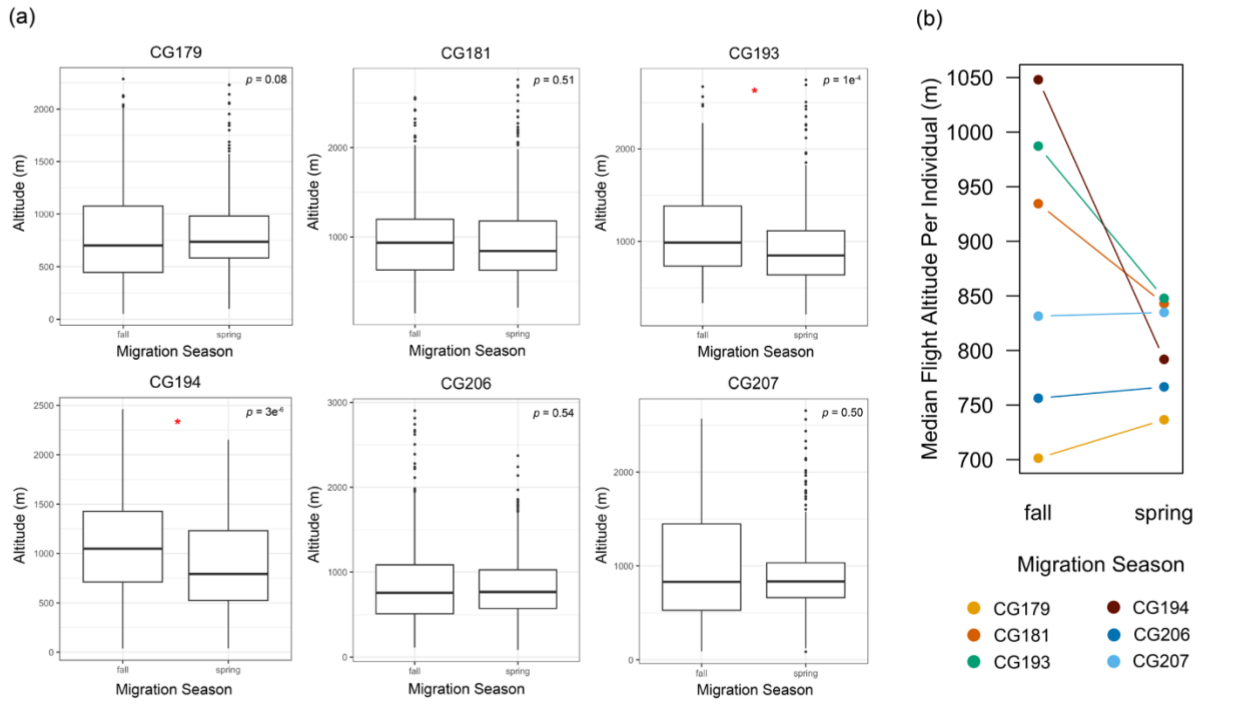


**Figure S23.** Flight altitude during fall and spring migration for six myrtle warblers determined from atmospheric pressure data collected by geolocators. (a) Boxplots showing distribution of flight altitudes in fall and spring migration for each bird. Pressure readings were recorded every 20 minutes to characterize changes in altitude over the course of a flight. Flight altitudes were significantly lower in spring compared to fall for two birds (indicated by red asterisks): CG193 (Wilcoxon rank sum test $W = 62832$, $p = 0.0001$) and CG194 ($W = 58425$, $p = 3e^{-6}$). (b) Interaction plot showing median flight altitude during fall vs. spring migration for each bird.

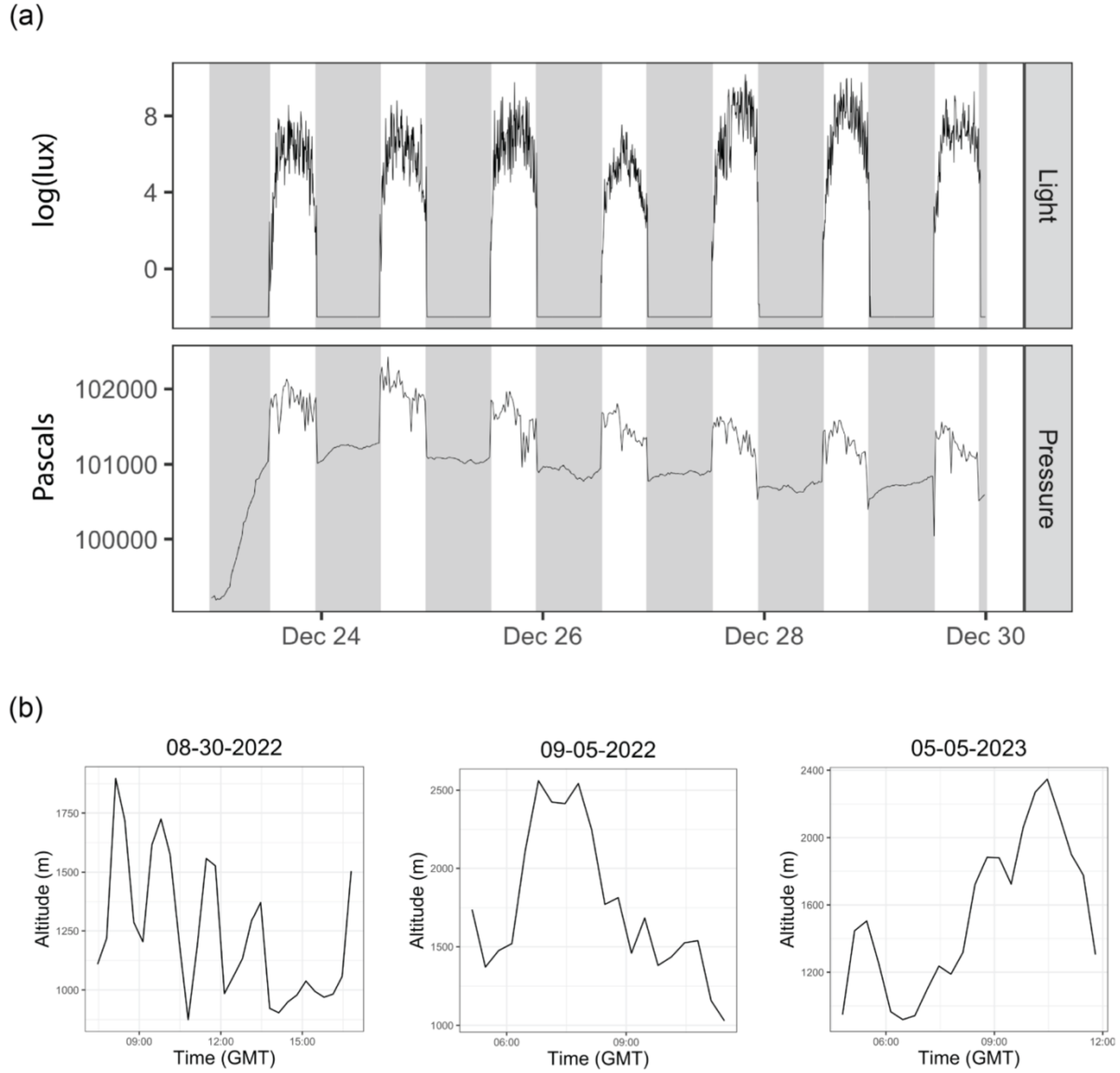


**Figure S24.** Fine scale changes in altitude observed from atmospheric pressure data collected by geolocators. (a) Over the wintering period, some birds exhibited cyclical changes in pressure that corresponded to periods of daylight and darkness, likely indicative of vertical movements related to foraging and roosting. (b) During a single flight, changes in pressure showed that birds flew at various altitudes throughout the flight.

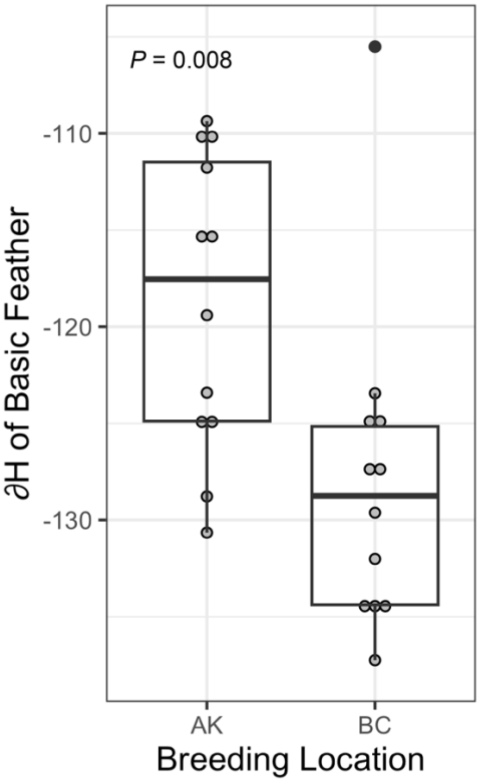


**Figure S25.** Comparison of stable hydrogen isotope values ($\partial^2$H) from basic feathers (likely grown on the previous year's breeding ground) of myrtle warblers breeding in Anchorage, AK vs. northern British Columbia. Feathers from birds breeding in British Columbia had significantly lower $\partial^2$H values than those from Alaska birds (W = 117, $p$ = 0.008).

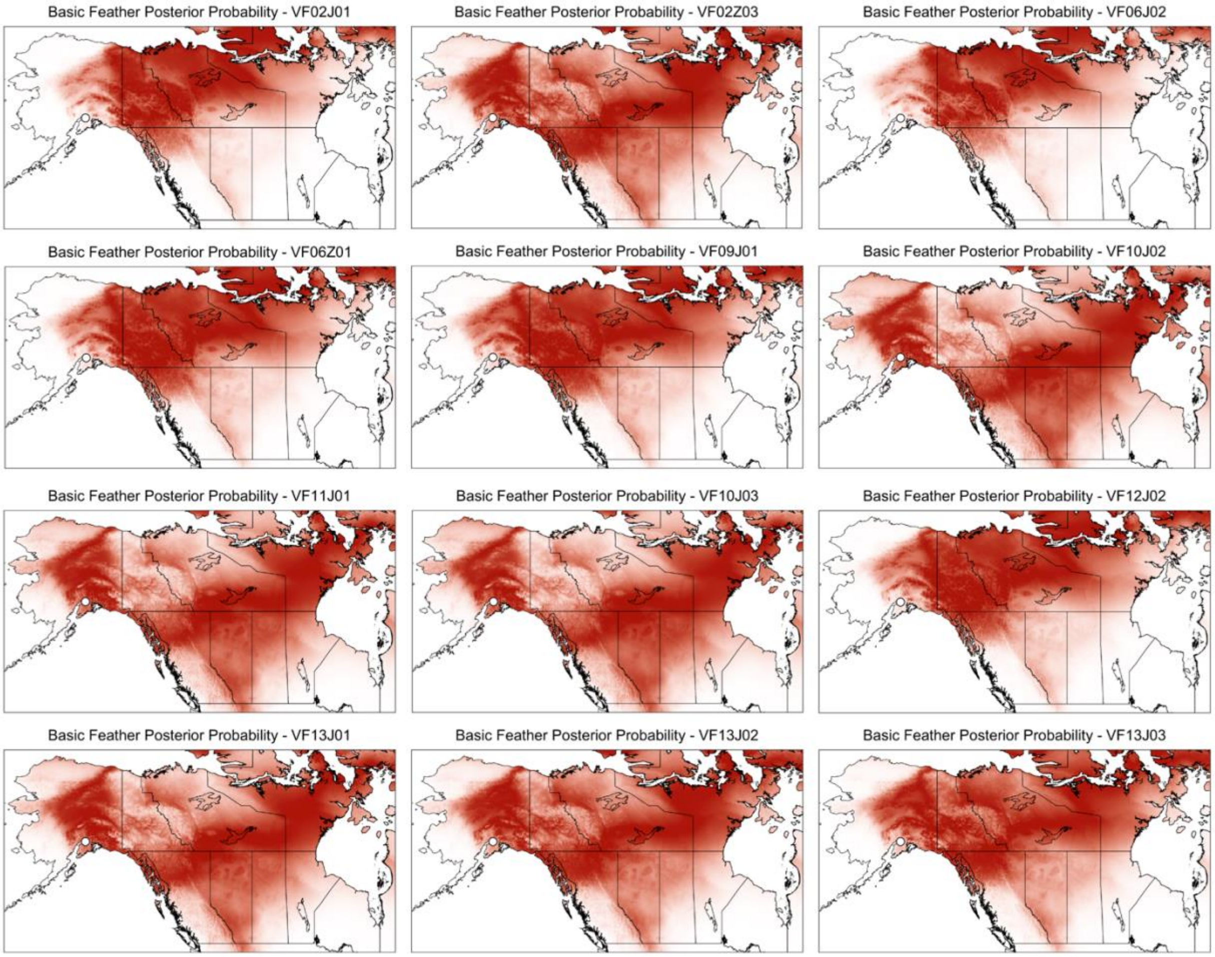


**Figure S26.** Stable hydrogen isotope posterior probability density maps for greater covert feathers grown in the pre-basic molt (likely on the previous year's breeding ground) of myrtle warblers breeding in Alaska. The white circle marks the sampling location, and darker red color indicates greater probability of origin.

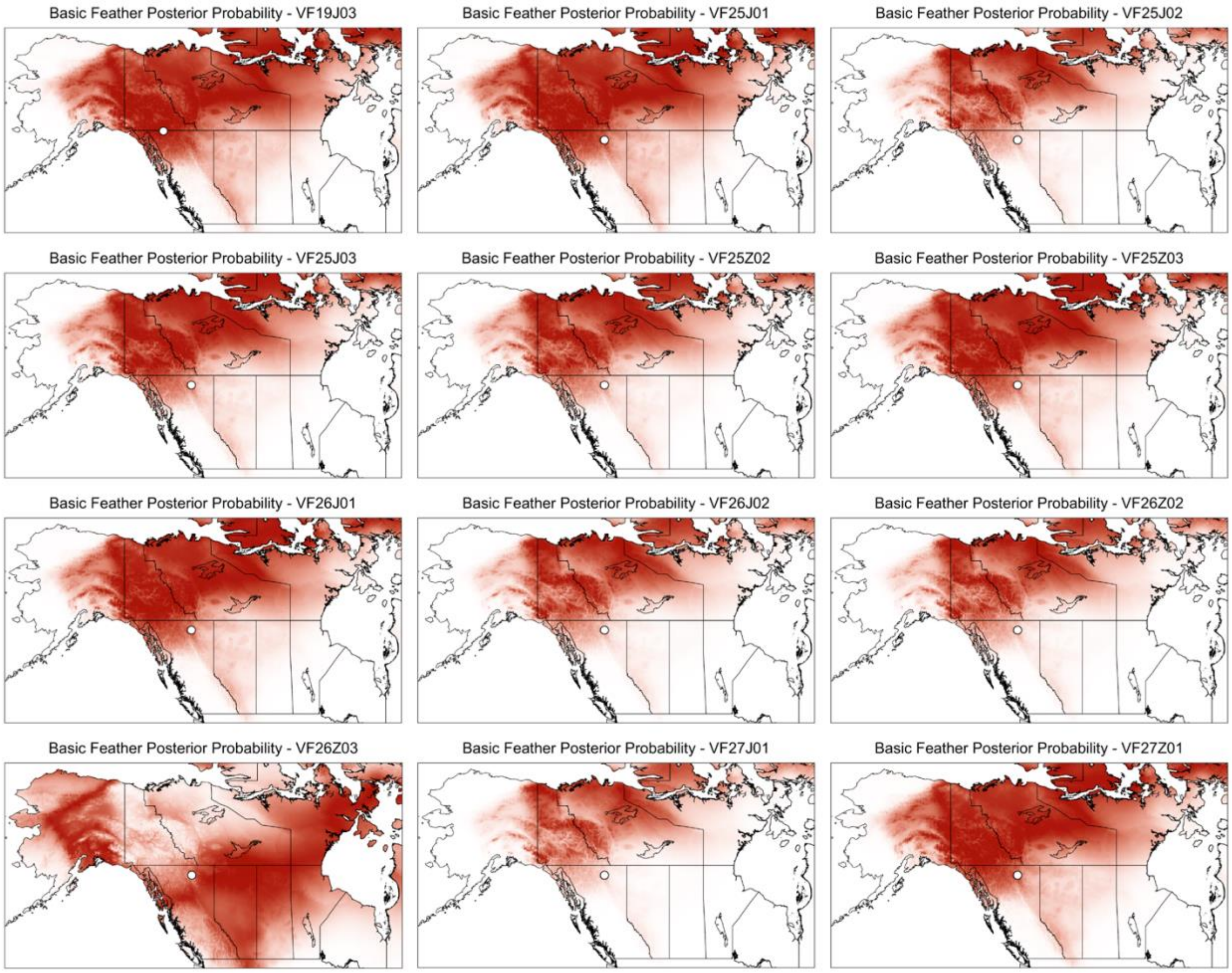


**Figure S27.** Stable hydrogen isotope posterior probability density maps for greater covert feathers grown in the pre-basic molt (likely on the previous year's breeding ground) of myrtle warblers breeding in northern British Columbia. The white circle marks the sampling location, and darker red color indicates greater probability of origin.

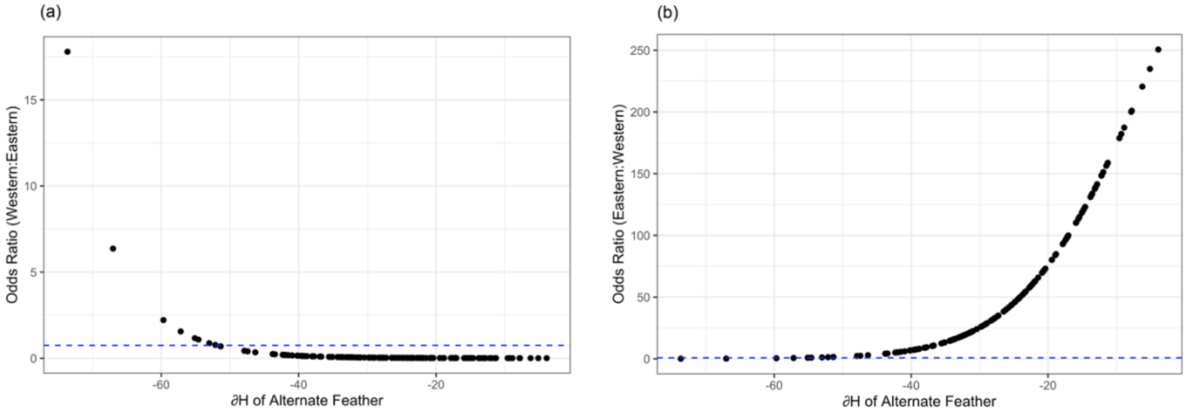


**Figure S28.** Odds ratios for wintering on the Pacific Coast ("western") vs. Gulf Coast ("eastern") nonbreeding areas for 167 myrtle warblers comparing posterior probabilities of origin for alternate covert feathers based on stable hydrogen isotopes. The dotted blue line indicates the ratio of the geographical areas of the two potential nonbreeding regions (0.75)—samples with an odds ratio of 0.75 are equally likely to have originated from either region. In (a) points above the blue dotted line represent feather samples with higher odds of originating from the western nonbreeding area, and in (b) points above the dotted line are samples with higher odds of originating from the eastern nonbreeding area.

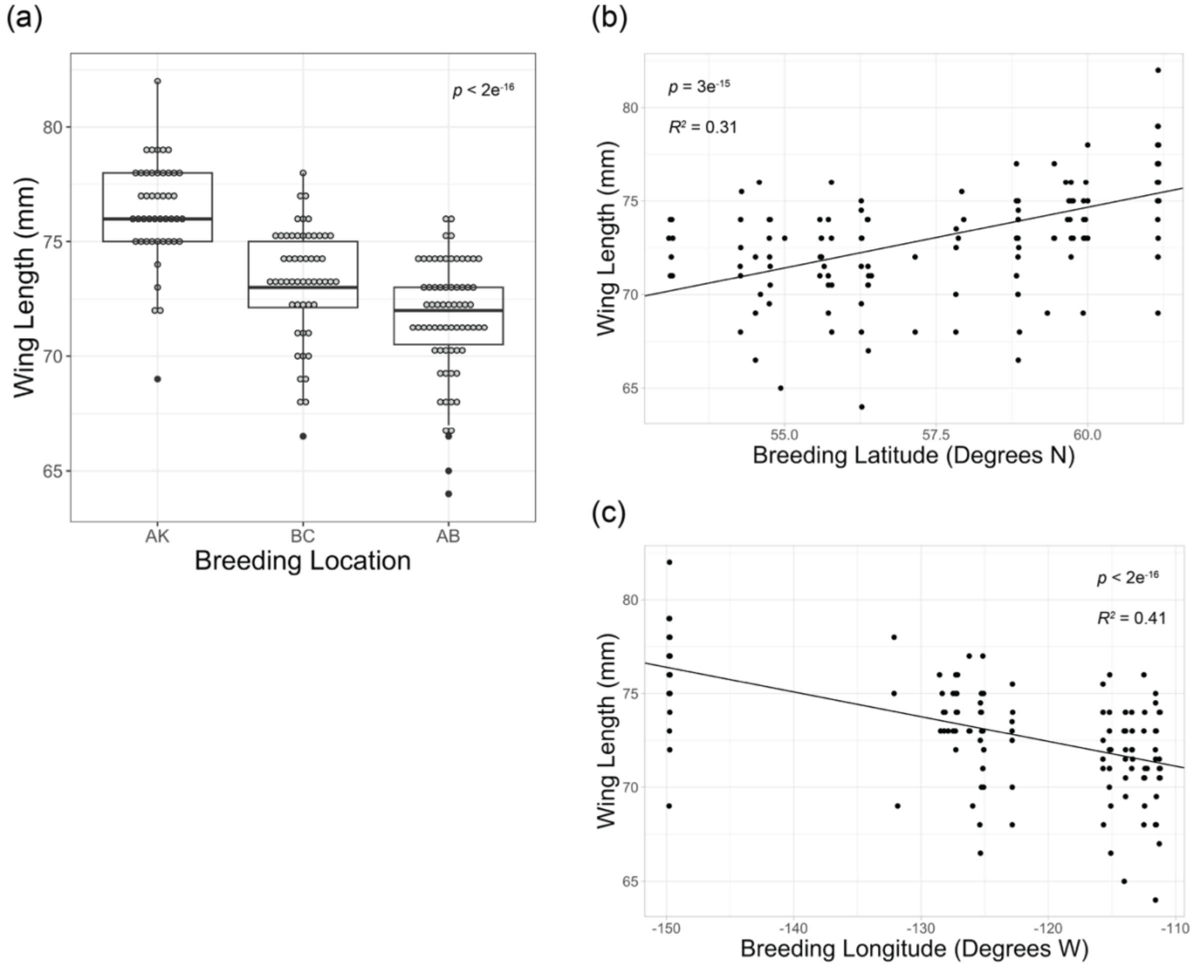


**Figure S29.** Relationship between wing length and breeding location for 167 myrtle warblers breeding in northwestern North America. (a) Wing length was significantly different between the three broad sampling areas: Alaska, British Columbia, and Alberta (Kruskal-Wallis $\chi^2$ = 73.418, *df* = 2, $p < 2.2e^{-16}$; Dunn's Test all pairwise comparisons $p <$ 0.008). (b) There was a significant positive relationship between wing length and breeding latitude ($p = 3e^{-15}$, $R^2 = 0.31$). (c) There was a significant negative relationship between wing length and breeding longitude ($p < 2e^{-16}$, $R^2 = 0.41$).

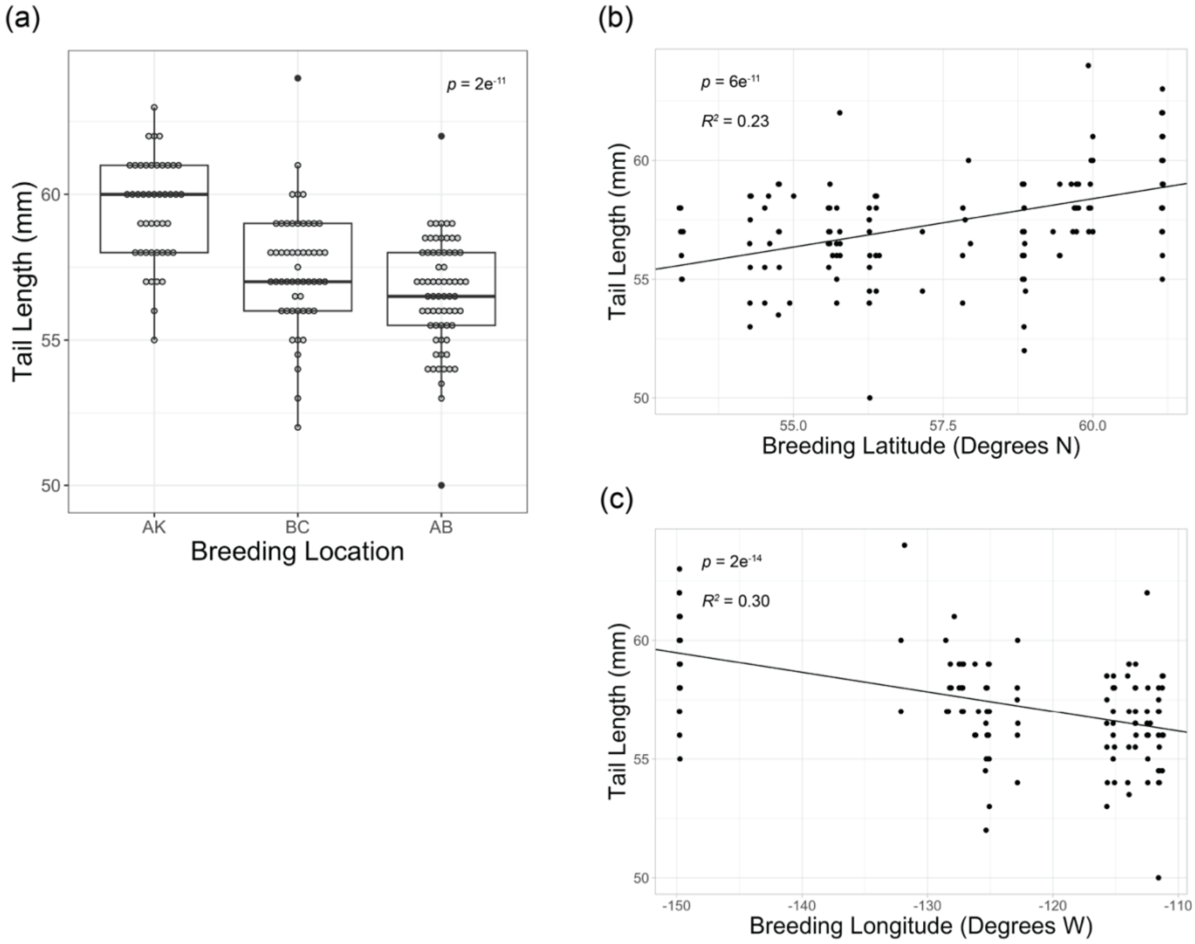


**Figure S30.** Relationship between tail length and breeding location for 167 myrtle warblers breeding in northwestern North America. (a) There was a significant difference in tail length among the three broad sampling areas (Kruskal-Wallis $\chi^2 = 49.527$, $df = 2$, $p = 1.8e^{-11}$). Tail length was significantly longer in Alaska than in British Columbia (Dunn's Test of pairwise comparisons $p = 7e^{-6}$) or Alberta ($p = 9e^{-12}$), but not significantly different between British Columbia and Alberta ($p = 0.11$). (b) There was a significant positive relationship between tail length and breeding latitude ($p = 6e^{-11}$, $R^2 = 0.23$). (c) There was a significant negative relationship between tail length and breeding longitude ($p = 2e^{-14}$, $R^2 = 0.30$).

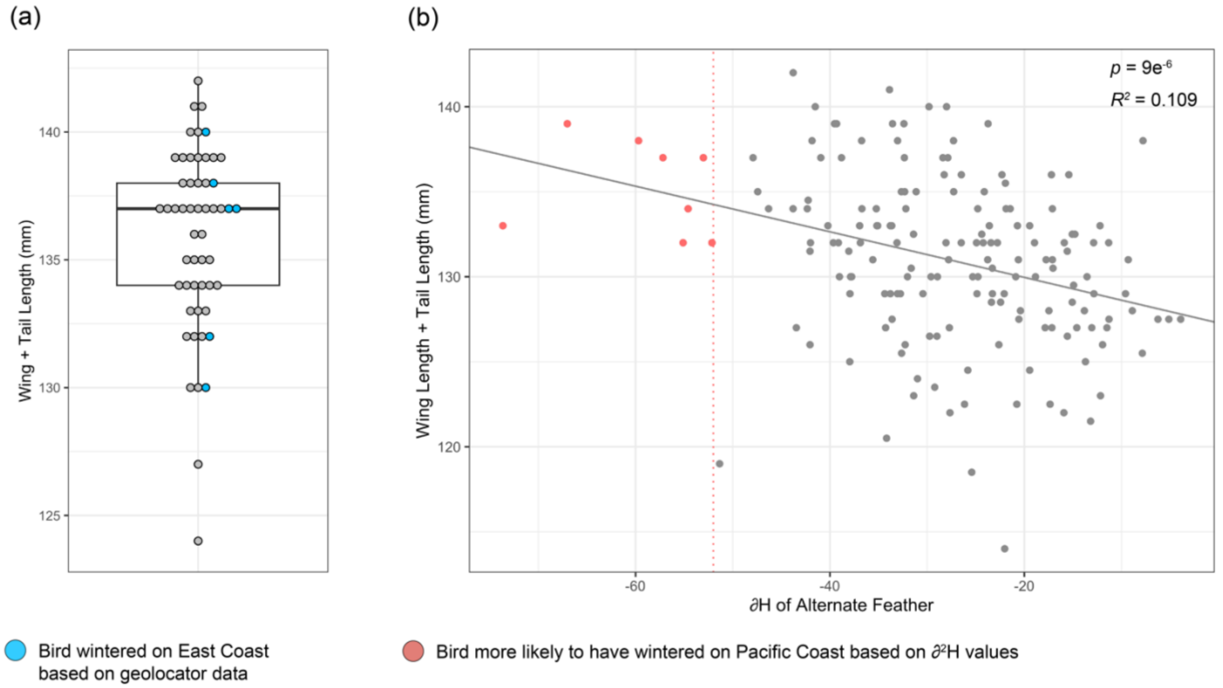


**Figure S31.** Relationships between wing and tail length and inferred wintering area. (a) Distribution of sizes (wing length + tail length) of myrtle warblers banded in Anchorage, AK. Blue points indicate the birds confirmed to breed on the Gulf Coast based on geolocator data. (b) Relationship between size (wing length + tail length) and stable hydrogen isotope ratio ($\partial^2$H) for all 167 myrtle warblers. All birds with $\partial^2$H values suggestive of Pacific Coast wintering (pink shading) had long wings and tails (i.e. "*hooveri*" type), but birds more likely to winter on the East Coast (grey) exhibited a range of sizes including many birds that historically would have been categorized as *S. c. hooveri*.

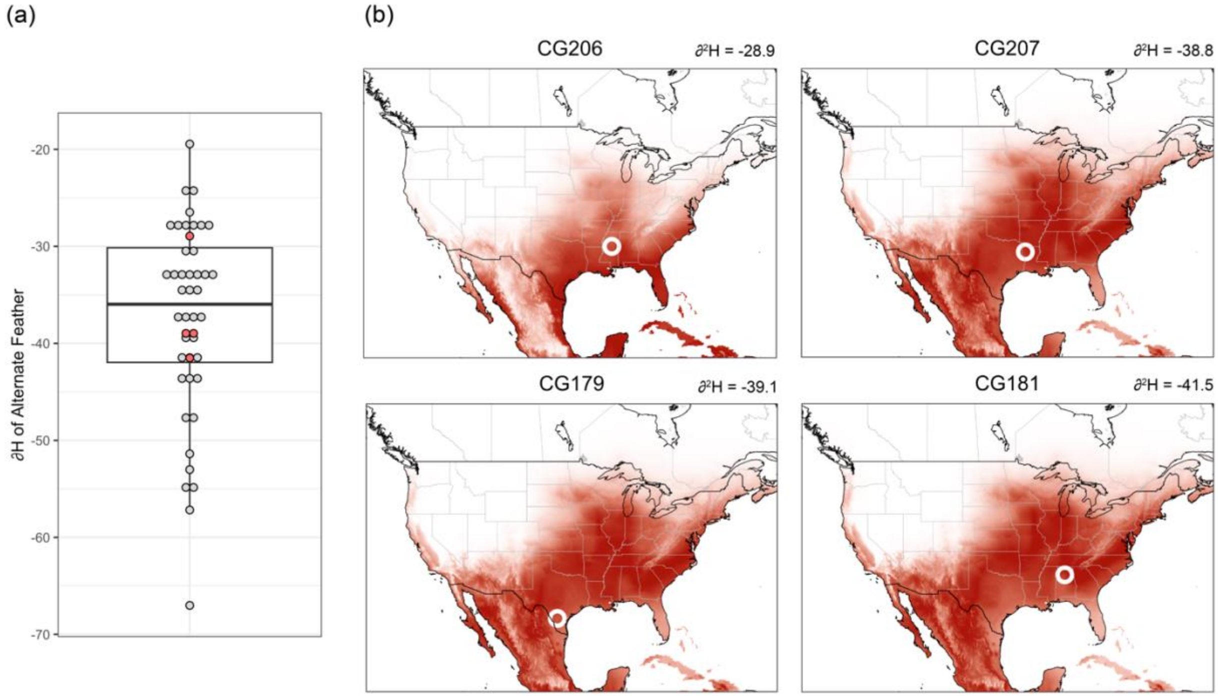


**Figure S32.** Stable hydrogen isotope ratios from feathers of myrtle warblers also tracked using geolocators. (a) Distribution of $\partial^2H$ for all birds sampled from Anchorage, AK with birds that also had geolocator data shaded in red. (b) Isotope likelihood surfaces for the four birds that had both isotope and geolocator data. Wintering areas inferred from geolocators are marked with a white ring.

### SUPPLEMENTARY TABLES

**Table S1.** Parameters used in estimation of twilight times and calibration of light-level geolocator data.

| Geolocator | Latitude of Deployment | Longitude of Deployment | Date of Deployment | Calibration Start Date | Calibration End Date | Light Threshold (lux) | Sun elevation angle (median) |
|---|---|---|---|---|---|---|---|
| CG179 | 61.154597 | -149.738660 | 3 June | 4 June | 25 June | 1.00 | -3.43 |
| CG181 | 61.158430 | -149.750052 | 2 June | 3 June | 24 June | 1.00 | -3.52 |
| CG193 | 61.165691 | -149.779153 | 14 June | 15 June | 6 July | 0.37 | -2.43 |
| CG194 | 61.162198 | -149.725350 | 7 June | 8 June | 29 June | 0.37 | -3.45 |
| CG206 | 61.157656 | -149.793144 | 13 June | 14 June | 5 July | 0.37 | -4.07 |
| CG207 | 61.159740 | -149.780571 | 11 June | 11 June | 2 July | 0.37 | -3.66 |

**Table S2.** Standard information for stable hydrogen isotope analysis at the Cornell University Stable Isotope Laboratory. Reported are mean and standard deviation $\partial^2H$ vs. VSMOW for three standards over three different runs.

| Run | CBS | | KHS | | Keratin | |
|---|---|---|---|---|---|---|
| | mean $\partial^2H$ | SD $\partial^2H$ | mean $\partial^2H$ | SD $\partial^2H$ | mean $\partial^2H$ | SD $\partial^2H$ |
| 1 | -156.95 | 2.43 | -35.35 | 1.04 | -47.30 | 2.19 |
| 2 | -156.90 | 2.88 | -35.40 | 2.50 | -47.26 | 2.83 |
| 3 | -156.92 | 2.68 | -35.38 | 2.14 | -49.68 | 2.49 |

**Table S3.** Point estimates for wintering areas of six myrtle warblers inferred using multi-sensor geolocators. Coordinates reported for GeoLight analysis are the average latitude and longitude of all point estimates generated using the threshold method over the longest winter stationary period. Coordinates generated using the GeoPressureR method represent the grid cell with the highest marginal probability for the trajectory model incorporating both light and pressure data. Also reported are the differences in latitude and longitude between the estimates from each method.

| | GeoLight | | GeoPressureR | | Latitude difference between methods | Longitude difference between methods |
|---|---|---|---|---|---|---|
| Geolocator | Latitude | Longitude | Latitude | Longitude | | |
| CG179 | 26.774 | -98.841 | 28.071 | - 98.929 | 1.30 | -0.09 |
| CG181 | 32.726 | -86.382 | 33.900 | -86.900 | 1.17 | -0.52 |
| CG193 | 35.132 | -81.241 | 35.083 | -81.917 | -0.05 | -0.68 |
| CG194 | 32.203 | -90.909 | 30.900 | -90.900 | -1.30 | 0.01 |
| CG206 | 37.632 | -89.008 | 33.100 | -89.500 | -4.53 | -0.49 |
| CG207 | 32.515 | -92.660 | 32.300 | -93.500 | -0.22 | -0.84 |

**Table S4.** Statistics describing migration timing for six myrtle warblers tracked using multi-sensor geolocators. Timing estimates were derived from atmospheric pressure data.

| Geolocator | CG179 | CG181 | CG193 | CG194 | CG206 | CG207 |
|---|---|---|---|---|---|---|
| *Fall migration* | | | | | | |
| Breeding site departure | 2 Sep | 29 Aug | 28 Aug | 5 Sep | 30 Aug | 4 Sep |
| Number of migratory flights | 18 | 16 | 19 | 19 | 19 | 19 |
| Average flight duration (hours) | 5.3 | 6.4 | 5.6 | 5.7 | 6.3 | 5.3 |
| Maximum flight duration (hours) | 8.7 | 10.7 | 10.0 | 11.7 | 13.0 | 13.7 |
| Average stopover length (days) | 5.2 | 4.1 | 3.2 | 4.5 | 4.7 | 3.9 |
| Total stopover time (days) | 89 | 61 | 58 | 80 | 81 | 70 |
| Wintering site arrival | 4 Dec | 2 Nov | 29 Oct | 28 Nov | 23 Nov | 17 Nov |
| Fall migration duration (days) | 93 | 65 | 62 | 84 | 85 | 74 |
| *Wintering period* | | | | | | |
| Days at wintering site | 111 | 158 | 158 | 89 | 127 | 133 |
| *Spring migration* | | | | | | |
| Wintering site departure | 26 March | 9 April | 5 April | 25 Feb | 30 March | 30 March |
| Number of migratory flights | 16 | 14 | 12 | 17 | 18 | 14 |
| Average flight duration (hours) | 7.6 | 8.1 | 9.4 | 6.1 | 8.9 | 7.6 |
| Maximum flight duration (hours) | 15.0 | 12.0 | 17.7 | 13.7 | 17.0 | 14.0 |
| Average stopover length (days) | 2.8 | 2.2 | 2.4 | 5.0 | 2.2 | 2.9 |
| Total stopover time (days) | 42 | 29 | 27 | 80 | 37 | 37 |
| Breeding site arrival | 11 May | 12 May | 6 May | 19 May | 12 May | 10 May |
| Spring migration duration (days) | 47 | 33 | 31 | 83 | 44 | 42 |
